# Discrete attractor dynamics underlying selective persistent activity in frontal cortex

**DOI:** 10.1101/203448

**Authors:** Hidehiko K. Inagaki, Lorenzo Fontolan, Sandro Romani, Karel Svoboda

## Abstract

Short-term memories link events separated in time, such as past sensation and future actions. Short-term memories are correlated with selective persistent activity, which can be maintained over seconds. In a delayed response task that requires short-term memory, neurons in mouse anterior lateral motor cortex (ALM) show persistent activity that instructs future actions. To elucidate the mechanisms underlying this persistent activity we combined intracellular and extracellular electrophysiology with optogenetic perturbations and network modeling. During the delay epoch, both membrane potential and population activity of ALM neurons funneled towards discrete endpoints related to specific movement directions. These endpoints were robust to transient shifts in ALM activity caused by optogenetic perturbations. Perturbations occasionally switched the population dynamics to the other endpoint, followed by incorrect actions. Our results are consistent with discrete attractor dynamics underlying short-term memory related to motor planning.

## Introduction

Short-term memory is the ability of the brain to maintain information over times of seconds. Neurons in the frontal cortex and related brain regions show persistent changes in spike rates during various types of short-term memory tasks ^1–15^. This persistent activity is a neural correlate of memory maintenance.

Short-term memory-related persistent activity has been extensively studied in delayed response tasks in non-human primates ^1, 4–6^. An instruction informs the type of action to be performed. A go cue determines the timing of action. The instruction and go cue are separated by a delay epoch, during which animals maintain a memory of the instruction and/or plan an upcoming movement. Persistent delay activity that predicts future actions is referred to as preparatory activity. Recently, preparatory activity has been observed in rodent models ^11–13, 16^ (Fig. 1a).

**Figure 1.**
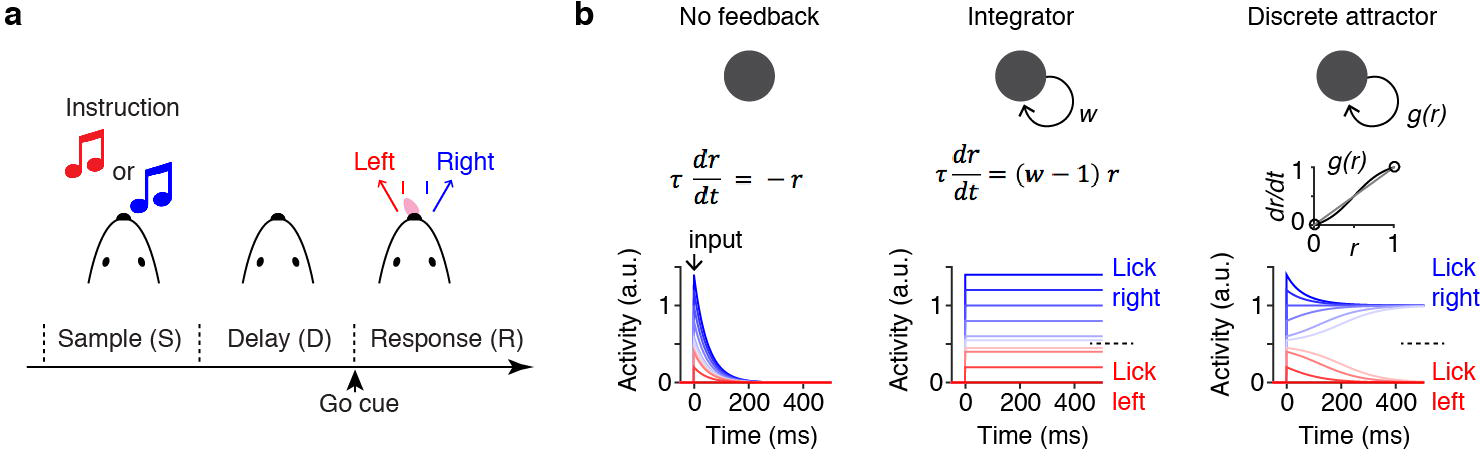
Models of persistent preparatory activity. **a.** Behavioral task. The instruction (tones for the auditory task, Fig 2–7; pole location for the tactile task, Fig 2–4) was presented during the sample epoch. The mouse reported its decision following the delay epoch by directional licking. The duration of the delay epoch was 1.2 (Fig.2–4) or 2.0 s (Fig.5, 6). **b.** Potential mechanisms underlying persistent activity. Left, in an isolated neuron, activity
caused by a brief input (arrow) decays following the membrane time constant of the cell. Middle, excitatory feedback can compensate for the decay to produce an integrator (*w* = 1), which has a continuum of stable states. Right, non-linear excitatory feedback, *g(r*) (inset), can convert the same circuit into one with two discrete attractors. Here activity has only two stable states.

In both tactile and auditory delayed response tasks, a large proportion of neurons in anterior lateral motor cortex (ALM) exhibit selective persistent activity, or preparatory activity, that predicts directional licking ^11,17^. Multiple lines of evidence indicate that preparatory activity in ALM is part of a multi-regional network mediating motor planning. First, unilateral inactivation of ALM during the delay epoch impairs future licking to the contralateral direction ^11,16^. Second, unilateral activation of pyramidal tract neurons in ALM during the delay epoch biases the future licking to the contralateral direction ^18^. Third, complete transient bilateral inactivation of ALM during the sample or delay epoch results in chance level performance ^16,19^, and loss of selective persistent activity ^19^. Yet, future licking directions can be decoded from population spiking activity in a trial-by-trial basis ^19^. The chance level performance after bilateral silencing during the sample epoch implies that other brain regions cannot rescue preparatory activity in ALM.

When current is injected into isolated neurons, activity decays within milliseconds, reflecting an interplay of rapid repolarizing currents and the neuronal membrane time constant ^20^ (Fig. 1b, left). A variety of theoretical models have been proposed to fill the gap in time scales between neuronal and network time constants ^7,21–38^. First, neurons might have specialized cell-autonomous mechanisms that maintain multi-stable persistent activity ^30,36,39–43^. Second, neurons could be wired into networks that effectively extend network time constants by sequential neuronal activation ^25,44,45^. Third, feedback excitation and inhibition ^23^ can compensate for dissipation of excitation. Depending on the structure of the circuit and the properties of individual neurons, the network can behave as an integrator with a continuum of stable (or quasi-stable) states ^23,31^, or a discrete attractor ^22,37,38^ (Fig. 1b, middle and right). Integrator and discrete attractor models can explain many aspects of neuronal activity and have been proposed to underlie short-term memory and decision making ^21–24,32–34,46^. Our previous analysis did not detect sequential activity in ALM during the delay epoch ^16^. Here we focus on distinguishing between other models of persistent activity, including cell-autonomous multi-stability, integrator networks and discrete attractor networks.

Based on dimensional reduction methods, population activity in ALM during the delay epoch can be mostly (more than 80 % of variance) explained by two modes ^47^. Such low-dimensional activity dynamics of short-term memory provides an opportunity to analyze neural activity and its response to optogenetic perturbations with little influence from non-mnemonic activity. Membrane potential measurements with whole cell recordings did not reveal evidence for cellular intrinsic multi-stability. We found that membrane potential and spiking dynamics funneled towards discrete endpoints and that these dynamics were robust to optogenetic perturbations. Occasionally, perturbations caused switches from one trajectory to the other, followed by incorrect choices. These data are inconsistent with integrator models, but are consistent with discrete attractor dynamics underlying short-term memory.

## Results

### The membrane potential underlying preparatory activity

We performed whole-cell recordings from left ALM (AP 2.5mm, LM 1.5mm, bregma) neurons during the delayed-response task (42 cells, tactile task; 37 cells, auditory task) (Fig. 2 and Extended Data Fig. 1, 2). Twenty of the recorded neurons were selective during the delay epoch (spike rate significantly different between correct lick-right (contra) and correct lick-left (ipsi) trials) (*ranksum test*, *p* < 0.05). Consistent with extracellular recordings ^11,16^, selectivity increased during the delay epoch (Fig. 2d, e). The membrane potential (Vm) to spike rate (SR) relationship was threshold linear ^48^ (Extended Data Fig. 1, 2). Even small and rapid features in the average Vm were reflected in parallel changes in SR (Extended Data Fig. 1, 2). The selectivity of Vm ramped during the delay epoch, similar to SR (Fig. 2d, e). Therefore, persistent changes in Vm underlie persistent changes in SR during the delay epoch.

**Figure 2.**
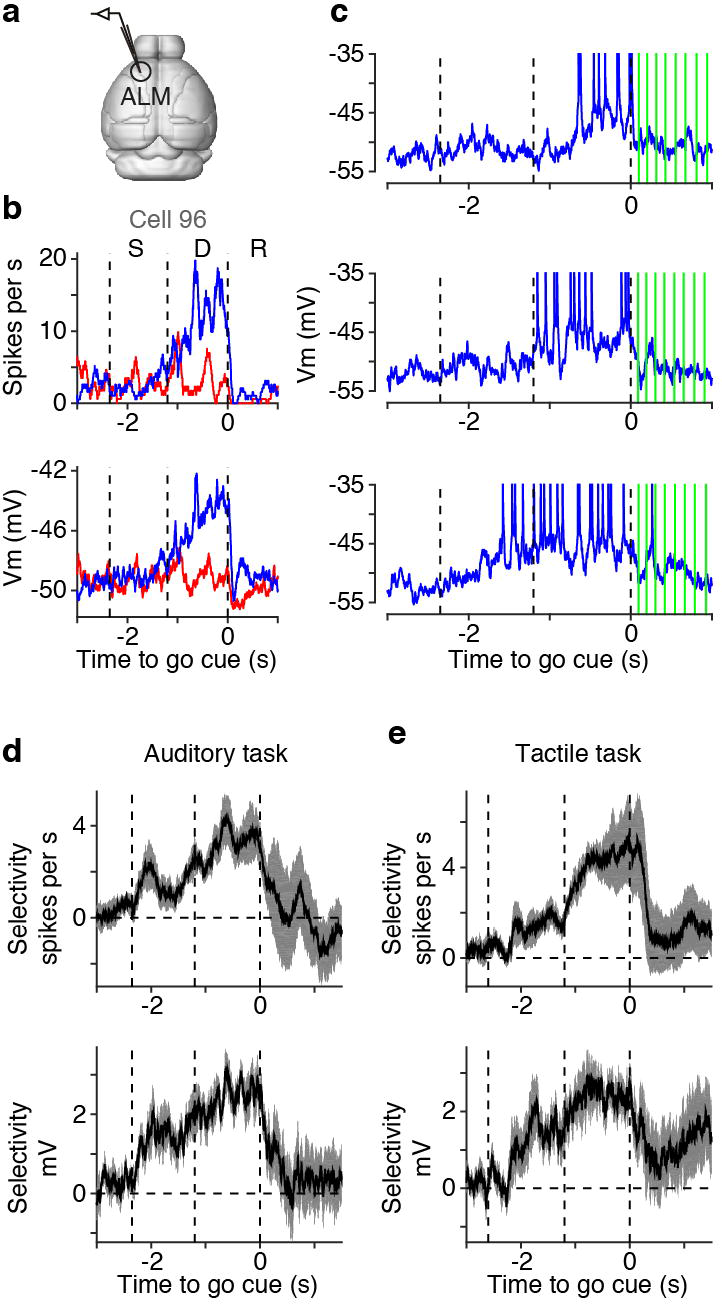
Whole cell recordings in ALM. **a.** Whole-cell recordings from left ALM. **b.** An example ALM neuron. Top, mean spike rate (SR); bottom, mean membrane potential (Vm). Blue, contra trials (correct lick-right trials); red, ipsi trials (correct lick-left trials). Time is aligned to the timing of the go cue. Dashed lines separate epochs. S, sample epoch; D, delay epoch; R, response epoch. **c.** Three example contra trials (same cell as b). Green lines, licks to the right lick port. **d.** Selectivity of ALM neurons based on SR (top) and Vm (bottom) in the auditory task. **e.** Same as **d**, tactile task.

### Testing for intracellular multi-stability

The membrane time constant limits how long isolated neurons can maintain activity after a transient input. The membrane time constant was short for selective cells (21.3 ± 19.7 ms, mean ± s.d., n = 19) and non-selective cells (20.5 ± 16.9 ms, mean ± s.d., n = 54). Moreover, the membrane fluctuations were faster in the delay epoch compared to the pre-sample epoch (Extended Data Fig. 3). Long membrane time constants therefore do not explain preparatory activity.

We next tested for other cell-autonomous mechanisms of multi-stability ^30,36,39–42^. First, spike bursts can activate voltage-dependent channels to trigger cell-autonomous persistent activity ^30,36,41,42^. However, only a small fraction of neurons (12/79 cells, 4/20 selective cells; > 1 complex spike / 5 trials) showed spike bursts (Extended Data Fig. 1c and 2c), and in these neurons spike bursts did not precede preparatory activity (Extended Data Fig. 1d and 2d). Therefore, spike bursts do not contribute to persistent activity in ALM.

Second, cell-autonomous mechanisms underlying persistent activity involve conductances activated by depolarization ^30,36,41,42^. Persistent activity should then be perturbed by hyperpolarization ^41^. For recordings with sufficiently long durations, we hyperpolarized cells by 12.0 ± 4.9 mV (mean ± s.d., n = 10). Hyperpolarized cells ceased to fire spikes but still showed selective changes in Vm (Fig. 3a, b and Extended Data Fig. 4a). Selectivity in Vm was similar after current injection; differences are most likely due to changing conductances (Extended Data Fig. 4b, c) and decreased inhibitory currents (hyperpolarization moved the membrane potential closer to the reversal potential for chloride). Although it is unlikely that we controlled membrane potential throughout the dendritic arbors ^49^, the fact that we silenced complex spikes, which require dendritic electrogenesis ^50^, with somatic hyperpolarization (Extended Data Fig. 4d) suggests that dendritic membrane potential was manipulated in some cases. In addition, cells with extremely low spike rates still showed selectivity in Vm (Extended Data Fig. 4e). These results indicate that spiking and conductances activated by depolarization are not necessary for selectivity in Vm. Altogether, cell-autonomous mechanisms do not explain persistent activity in ALM ^51^. Instead, spike rate changes and membrane potential changes during the delay epoch were most likely driven by synaptic input.

**Figure 3.**
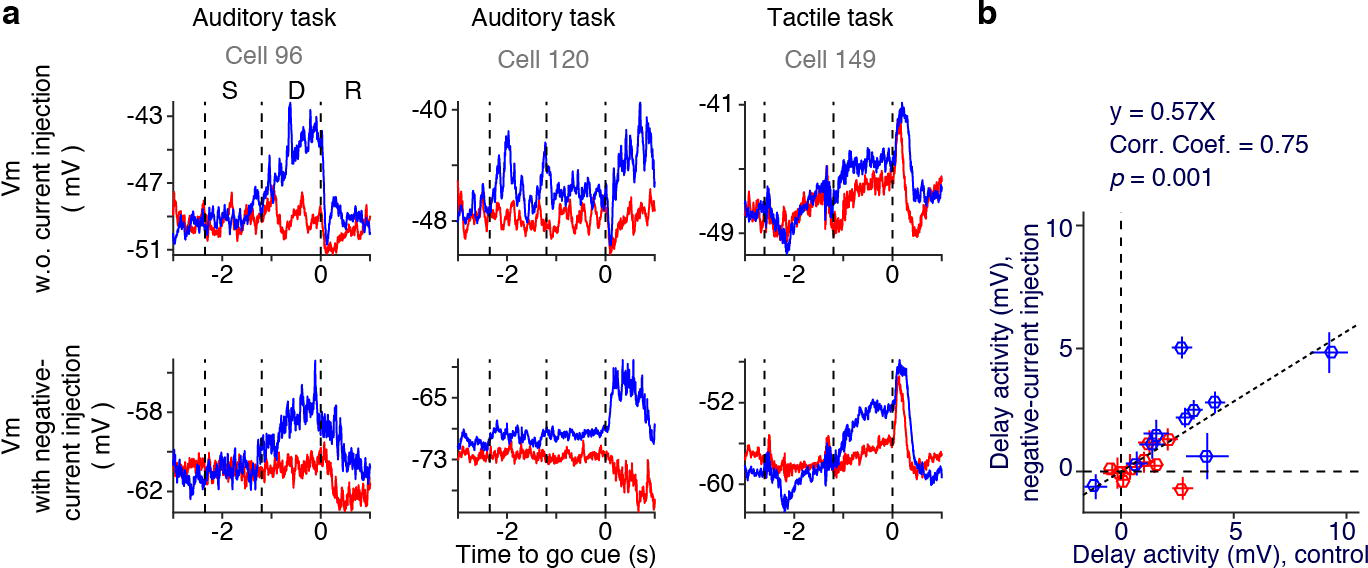
Hyperpolarization of ALM neurons. **a.** Example ALM neurons. Top, mean Vm without current injection; bottom, mean Vm with negative current injection. **b.** Delay activity of Vm (increase during the delay epoch relative to the baseline pre-sample epoch) with and without current injections (*n* = 10). Contra trials (blue) and ipsi trials (red) are shown separately. Crosses, S.E.M. (*bootstrap*). Dashed line, linear regression. Slope of linear regression, *Pearson* correlation coefficient (Corr. Coef.), and the *t-statistic* of *Pearson* correlation coefficient (*p*) are shown.

### Funneling of membrane potential

We now focus on network mechanisms (Fig. 1b). In a system following discrete attractor dynamics, activity is expected to converge to discrete endpoints over time (funneling). In contrast, in a system following integrator dynamics, funneling is not expected (Fig. 1b). During the delay epoch, Vm funneled to a narrow distribution at the time of movement onset (Fig. 4a, Extended Data Fig. 5a). We quantified funneling by computing the across-trial fluctuations in Vm (the difference between the first and third quartiles of Vm across trials at each time point). Across-trial Vm fluctuations decreased during the delay epoch, and reached a minimum immediately after the go cue (Fig. 4b, Extended Data Fig. 5d, f). This decrease was strong in contra trials but weak in ipsi trials (Fig. 4c, d, Extended Data Fig. 5) (*Hierarchical bootstrap* comparing across-trial fluctuations in the baseline and the delay epoch, Methods). The reduction in Vm fluctuations was stronger in selective neurons (Fig.4c, d, Extended Data Fig. 5d-g).

**Figure 4.**
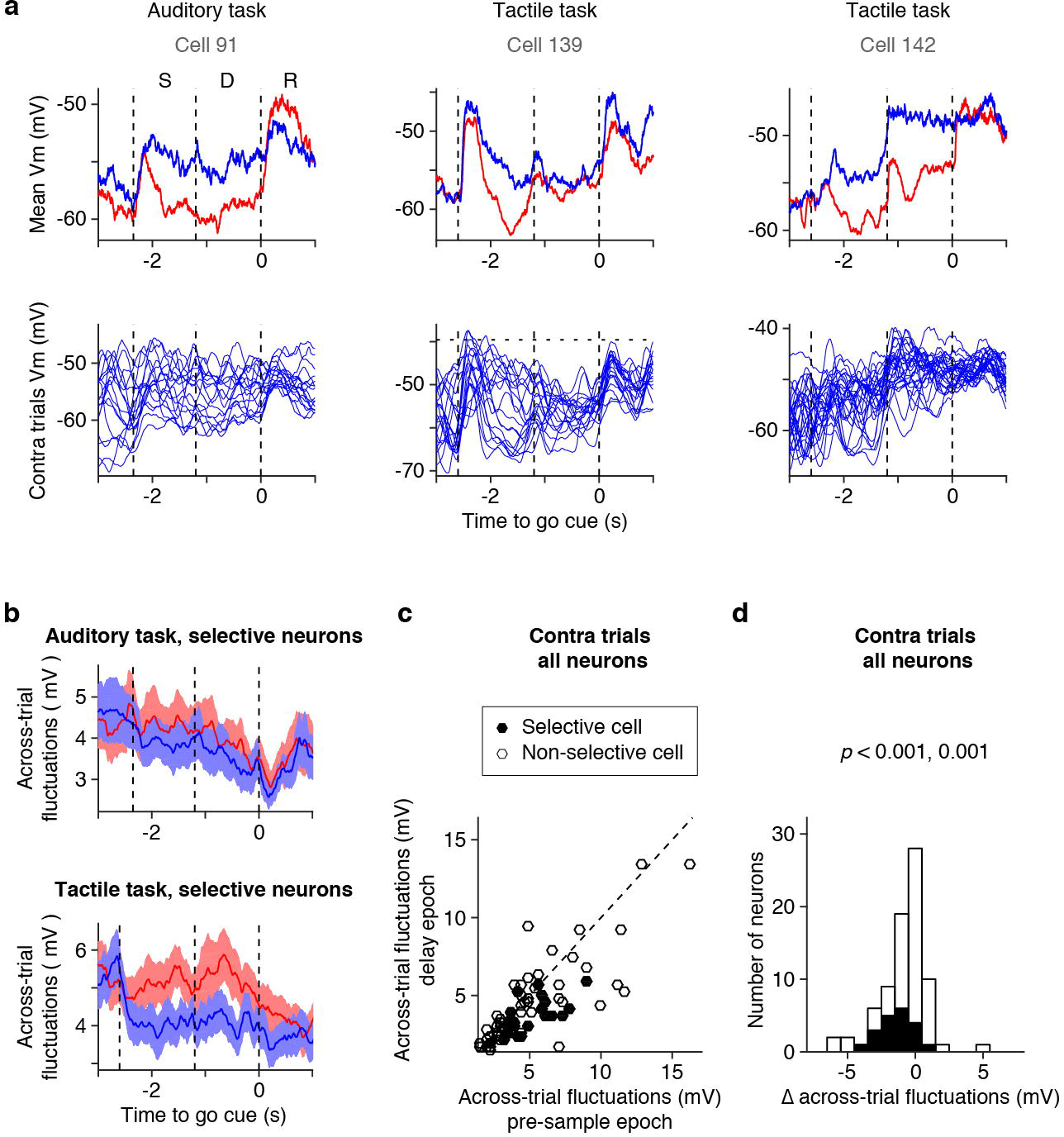
Funneling of Vm. **a.** Example ALM neurons. Top, mean Vm; Bottom, all contra trials overlaid. Vm was averaged over 200 ms. Horizontal dashed line (Cell 139), spike thresholds. In other cells, spike thresholds were higher than the range shown in the figure. **b.** Across-trial fluctuations of selective cells in the auditory task (top, *n* = 10 cells) and in the tactile task (bottom, *n* = 10 cells). Line, mean of across-trial fluctuations among cells; Shadow, S.E.M. (*hierarchical bootstrap*, Methods). Across-trial fluctuations are the difference between first and third quartile of all trials within the same trial type. **c.** Relationship between across-trial fluctuations during the pre-sample epoch and during the delay epoch in the contra trials. For **c** and **d**, *n* = 79 cells; Black, selective cells (*n* = 20 cells). **d.** Distribution of difference in across-trial fluctuations between the pre-sample epoch and the delay epoch (Δ across-trial fluc.). *P*-values, *signrank* test examining a null hypothesis that Δ across-trial fluc is 0. The first *p*-value, all cells (2.8×10^−5^); second *p*-value, selective cells only (5.2×10^−4^).

The reduction in across-trial Vm fluctuations was not caused by a ceiling effect imposed by the spike threshold. First, the threshold linear Vm-to-SR relationship (Extended Data Fig. 1 and 2) indicates that the activity during the delay epoch was not close to saturation. Second, the reduction in across-trial Vm fluctuations was independent of the distance to spike threshold (Extended Data Fig. 5h).

Altogether we found that the across-trial Vm fluctuations decayed during the delay epoch in contra trials, especially in selective cells, consistent with funneling. A reduction in across-trial variability in spike count has been reported during motor planning in non-human primates ^52,53^. However, the interpretation of fluctuations based on spike counts is complex because both the across-trial variance of SR and the variance of the point process contribute to the overall variance ^54^. Our whole-cell recording data show a reduction in variability of Vm, which controls SR.

### Funneling of delay activity

The attractor model predicts funneling of population activity along dimensions that predict movement direction. We analyzed populations of ALM neurons recorded simultaneously using high-density silicon probes during the auditory task (28.2 ± 12.4, mean ± s.d., putative pyramidal cells per session, from 6 animals, 20 sessions). The activity of *n* neurons describes an *n*-dimensional activity space. In this space we refer to the direction that best predicts the future action (i.e. right or left lick) as the coding direction (CD)^19^ (Extended Data Fig. 6a) (Methods).

For each recording session, we projected population activity of individual trials to the CD (Fig. 5a) (Methods). We observed large variability in the projected population activity (Fig. 5a, Extended Data Fig. 6b). Since we were interested in variance related to fluctuations in the SR, but not in the point-process noise, we averaged trajectories as follows: trajectories were rank-ordered based on their activity 1.3 seconds before the go cue and 15 neighboring trajectories were averaged. The averaged trajectories reflect SR dynamics starting from a similar activity level. Averaged trajectories funneled toward discrete endpoints, depending on the trial type (Fig. 5b). Across-trial fluctuations of the trajectories decreased during the delay epoch (Fig. 5d, Extended Data Fig. 6c). This reduction was bigger in the contra than in the ipsi trials, consistent with the Vm measurements (Fig. 5d, e).

**Figure 5.**
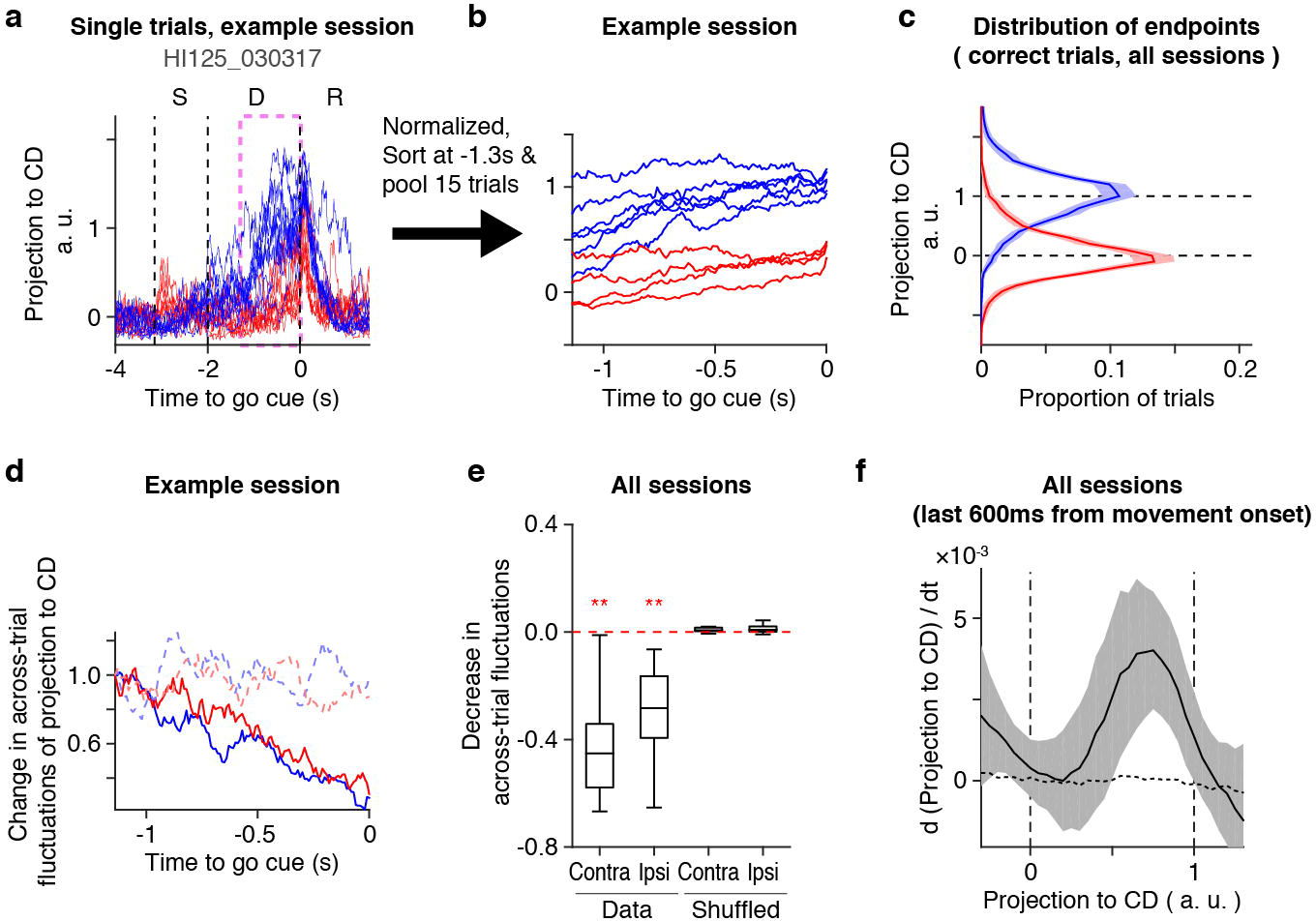
Funneling of activity trajectories along the coding direction. **a.** Projection of trials to coding direction (CD) in an example session (HI125_030317). Projections were normalized so that the medians of contra and ipsi distributions correspond to 1 and 0, respectively (See **c**.). Randomly selected 10 trials are overlaid. The pink box shows the time window analyzed in **b** and **d**. **b.** Trajectories in the example session. Trajectories (as in **a**) were rank ordered based on activity −1.3 s before the go cue and 15 adjacent trials were averaged. **c.** Distribution of endpoints (activity level in the last bin before the go cue; Methods). Mean of all sessions. Shading, S.E.M. (*hierarchical bootstrap*). Distribution of contra trials (blue), and ipsi trials (red) are shown. **d.** Across-trial fluctuations of the pooled trajectories in the example session (same as **a**, **b**). Across-trial fluctuations are normalized to the value at −1.2 s before the go cue. Solid line, data; dashed line, shuffled data (Methods). **e.** Difference of across-trial fluctuations between time 0 and −1.2. Central line in the box plot indicates median. Top and bottom edges are 75 % and 25 % points, respectively. The whiskers show the lowest datum still within 1.5 interquartile range (IQR) of the lower quartile, and the highest datum still within 1.5 IQR of the upper quartile. **: *p* < 0.01 in *one-sided signrank test* with *Bonferroni* correction, testing a null hypothesis that decrease in fluctuations is 0. From left to right *p* = 4.8×10^−5^, 1.1×10^−4^, *p* > 0.99, 0.99 (*p*-values without *Bonferroni* correction). **f.** Phase portrait of trajectories along CD, Solid line, data; dashed line, shuffled data; shading, S.E.M. (*hierarchical bootstrap*). The non-linearity disappeared in trajectories based on shuffled data.

The distribution of the endpoints just before the go cue was bimodal (Fig. 5c, Extended Data Fig. 6d, e). One peak corresponds to the contra trials, and the other to ipsi trials. Therefore, activity along the CD funneled toward discrete endpoints.

We next analyzed how the activity along CD relates to the drift of trajectories along the CD ^55^. For integrators, drift should be independent of the activity level. For discrete attractor models drift should be larger further from the endpoints, and near zero close to the endpoints (Projection to CD near 0 or 1). The negative slopes around the endpoints indicate stable fixed points (Fig. 5f, Extended Data Fig. 6f)^55^. Furthermore, the slope around the contra endpoint was sharper compared to the ipsi endpoint (Fig. 5f, Extended Data Fig. 6f). This indicates stronger attraction of ALM dynamics to the contra endpoint, consistent with the stronger funneling in contra trials (Fig. 5b). This dynamic behavior is consistent with discrete attractor dynamics (Extended Data Fig. 7).

### Robustness to perturbations

Funneling of activity along the CD is consistent with discrete attractor dynamics. However, it is possible that this funneling reflects dynamics outside of ALM ^56^ (Extended Data Fig. 7I). To further distinguish between integrator and attractor models, we transiently distorted neural trajectories by applying optogenetic manipulation in the middle of the delay epoch (Fig. 6a). If ALM preparatory activity evolves with discrete attractor dynamics, activity should still funnel toward one of the discrete endpoints after the perturbation (Fig. 6b, c and Extended Data Fig. 7).

**Figure 6.**
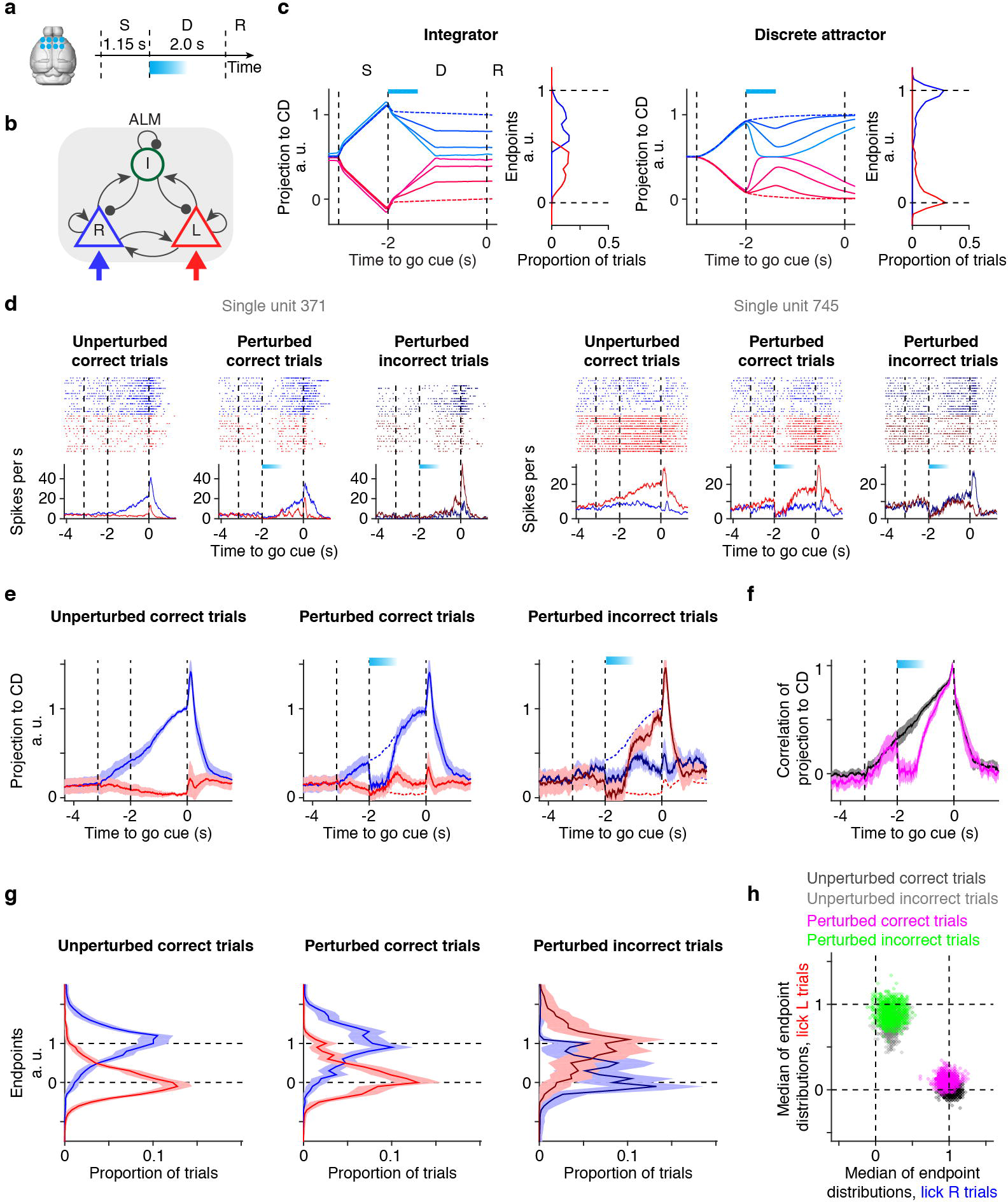
Robustness of discrete trajectories. **a.** Schematics of experiments. ALM was photoinhibited bilaterally during the first 600 ms of the delay epoch with 400 ms ramp down. **b.** Circuit model (related to **c**). R, L, and I correspond to lick right selective pyramidal neurons, lick left selective excitatory neurons, and inhibitory interneurons, respectively. Blue and red arrows, selective external input. **c.** Expected behavior of integrator (left) and discrete attractor (right) (Methods). **d.** Example single units. Top, spike raster. Fifteen trials per trial type were randomly selected. Bottom, mean SR. Blue, contra trials; red, ipsi trials; dark blue, incorrect lick right trials; dark red, incorrect lick left trials. Blue box, photoinhibition. **e.** Trajectories along CD. Grand mean trajectories of unperturbed correct trials (left). Grand mean trajectories of perturbed trials followed by correct lick (middle) or incorrect lick (right) are overlaid on top of trajectories of unperturbed correct trials (dashed lines). Trials from all sessions were pooled. Line, mean; shading, S.E.M. (*hierarchical bootstrap*). **f.** Pearson correlation of CD trajectories at each time point to that at the endpoint. Black, unperturbed correct trials; magenta, perturbed correct trials. Both contra and ipsi trials were pooled. **g.** Distribution of endpoints in e. Shading, S.E.M. (*hierarchical bootstrap*). Distributions of contra trials (blue), and ipsi trials (red) are shown. **h.** Median of the endpoint distributions in **g**. X-axis, Median of endpoint distributions in lick right trials (blue in **g**); Y-axis, Median of endpoint distributions in lick left trials (red in **g**). Black, unperturbed correct trials (left in **g**); Grey, unperturbed incorrect trials; Magenta, perturbed correct trials (middle in **g**); Green, perturbed incorrect trials (right in **g**). Each dot represents one iteration of *hierarchical bootstrap*.

We activated GABAergic neurons in mice expressing ChR2 in fast-spiking interneurons (photoinhibition, Methods). We performed bilateral photoinhibition of ALM during the first 0.6 s (with additional 0.4 s of ramping down) of the delay epoch. Consistent with previous work ^16,19^, photoinhibition with strong laser power (1.5 mW per spot; median activity 2.4 % of baseline) resulted in near chance level performance (Extended Data Fig. 8a, b), confirming that ALM is required for the short-term memory. For perturbations, we used more modest photoinhibition (median activity 20 ~ 60 % of baseline; Extended Data Fig. 8 and 9a, b). Under these conditions rebound spiking ^11^ was small and did not cause early behavioral responses.

In perturbed trials that resulted in correct movement, selectivity recovered (784 ± 259 ms, mean ± s.e.m.) to the trajectories of unperturbed trials on average (Fig. 6d and Extended Data Fig. 9b-d). In trials with incorrect movement direction, these neurons switched trajectories to that of the other trial type (Fig. 6d and Extended Data Fig. 9d). This switching effect was more prominent in preparatory cells, which predict future licking direction (Extended Data Fig. 9d top), compared to other cells (Extended Data Fig. 9d bottom). Therefore, the activity trajectories snap to one of the two discrete endpoints, independent of behavioral outcome (correct or incorrect).

Similar to individual neurons, trajectories along CD recovered in perturbed trials followed by correct movement (Fig. 6e, middle). The *Pearson* correlation of CD trajectories at each time point to that at the endpoint showed recovery of trajectories after the perturbation (Fig. 6f). Furthermore, we found that CD trajectories moved toward the opposite endpoint in perturbed trials followed by incorrect lick (Fig. 6e, right). Endpoints of activity along CD were discrete for both perturbed and unperturbed trials with peaks at identical locations (Fig. 6g, h and Extended Data Fig. 9e-g). The convergence to endpoints was weaker for lick left trials (correct ipsi trials and incorrect contra trials; red in Fig. 6e middle, and blue in Fig. 6e right), consistent with weaker attraction to this endpoint (Fig. 5f).

Since animals made errors even without perturbation (Performance, 87.0 ± 8.4 %, mean ± s.d.), some of the perturbed trials may reflect errors independent of the perturbation. We decoded the future licking direction based on activity before the perturbation (Methods). Trials decoded to be correct before the perturbation (performance of decoder, 77.2 ± 16.2 %, mean ± s.e.m.) (Extended Data Fig. 9h) showed switching in the endpoints in incorrect trials (Extended Data Fig. 9i, j), implying that switches were caused by the perturbation. Similarly, we observed trials that were decoded to be incorrect before the perturbation and then switched trajectories after perturbation to become correct (Extended Data Fig. 9k, l). The recovery and switching of CD trajectories toward discrete endpoints after perturbation is consistent with discrete attractor dynamics in ALM.

### Relationship between discrete attractor dynamics and ramping activity

In a variety of behavioral tasks and species, preparatory activity ramps up to a movement ^4––13,15,16^ (Fig. 6). In contrast, discrete attractor models show stationary activity once the fixed points are reached. Ramping dynamics with discrete attractor networks either requires tuning of network parameters to generate slow decay to the fixed points^19^, or a non-selective ramping input that moves fixed points apart over time (Extended Data Fig. 11a-i vs. j-q).

We explored these possibilities further. Ramping predicts the timing of movement^10,57,58^. We performed a separate set of recordings with randomly selected delay durations (Fig. 7a, b and Extended Data Fig. 10a, b) (302 units, 11 sessions, 4 animals), which precludes prediction of the timing of movement. Similar to the fixed delay task^16^, many neurons (99 out of 260 pyramidal neurons) showed selective persistent activity during the delay epoch.

**Figure 7.**
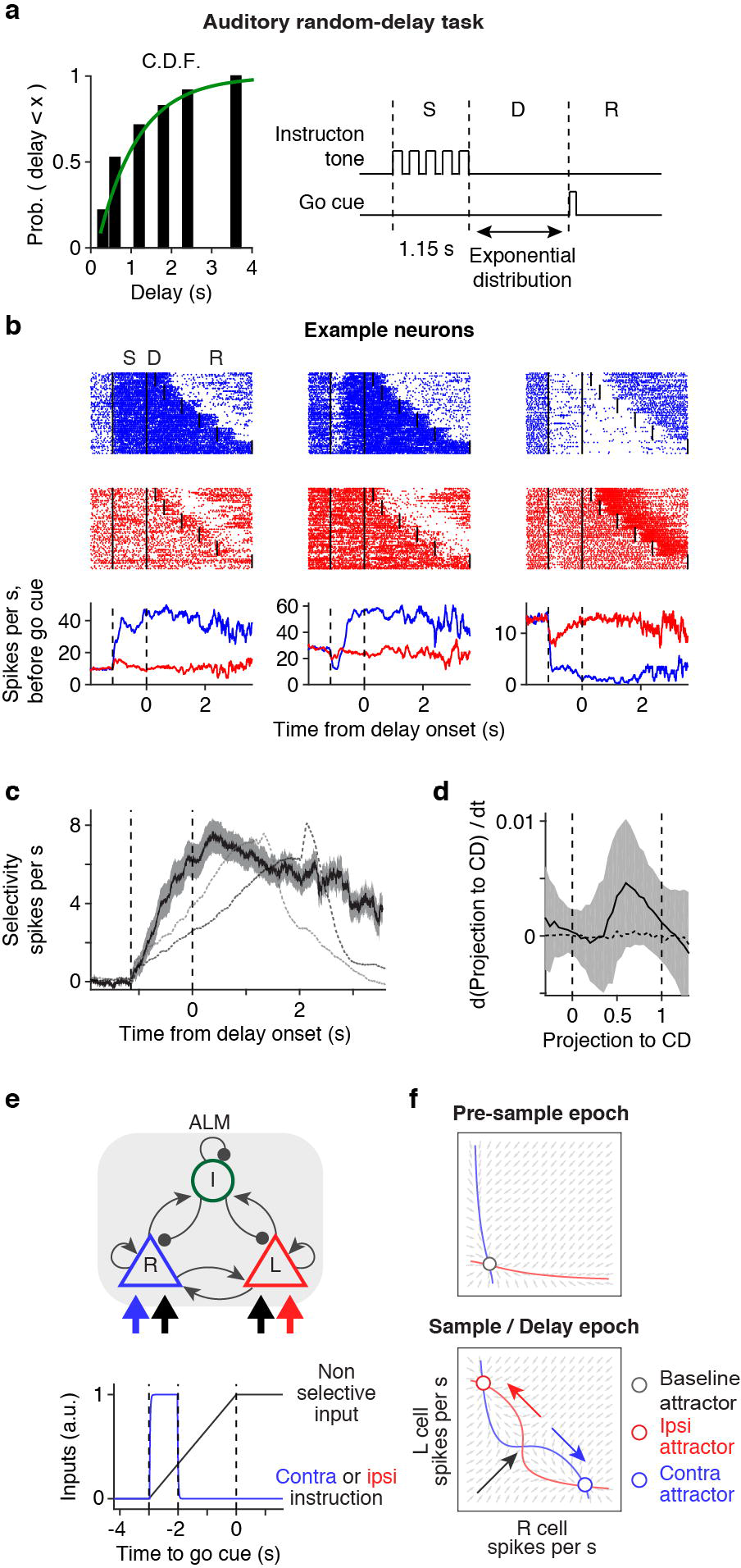
Stationary preparatory activity. **a.** Delays in each trial were randomly selected from six durations (0.3, 0.6, 1.2, 1.8, 2.4, 3.6 s). The cumulative distribution function (C.D.F.) of delay durations (black bar) approximated the C.D.F. of the exponential distribution (green line). **b.** Example cells. Top, spike raster. Lines separate epochs. Ten trials per delay duration were randomly selected. Bottom, mean SR pooling spike before the go cue across trials with different delay durations. Because delay durations were different across trials, the time axis was aligned to the onset of the delay epoch (same in **c** and **d**). **c.** Selectivity. Solid line, mean selectivity in random delay task. Shading, S.E.M. (*bootstrap*). Dashed lines, selectivity in fixed delay tasks (grey, 1.2 s; black, 2.0 s. Data from ^47^). **d.** Phase portrait of trajectories along CD. Solid line, data; dashed line, shuffled data; shading, S.E.M. (*hierarchical bootstrap*). **e.** Model schematics. Top, circuit architecture. Bottom, timing of external input. Black arrows, non-selective external input. **f.** Phase portrait of the model. Blue and red lines, nullclines of lick right and left populations, respectively. Black, blue, and red arrows indicate the effect of non-selective input, contra instruction and ipsi instruction, respectively.

Spike rates and selectivity ramped up rapidly, before the first possible go cue, and then remained near stationary during the delay epoch (Fig. 7b, c and Extended Data Fig. 10c, d). The CD and non-selective mode together explained 81.1 ± 11.8 % (mean ± s.e.m) of variability in activity during the delay epoch (Methods). Neither mode shows ramping up during the delay epoch, unlike in the fixed delay task (Extended Data Fig. 10e-i). To examine the possible contribution of ramping activity to the reduction in across-trial fluctuations, discreteness of endpoint distributions, and CD phase portrait, we repeated these analyses on data from the random delay task (Fig. 7d and Extended Data Fig. 10j, k). In all cases, the results are similar to those observed in the fixed delay task. These analyses confirmed that CD in the random delay task still followed discrete attractor dynamics.

Previous studies have shown that the non-selective mode recovers to ramp even after bilateral silencing of ALM, unlike activity along the CD ^19^. The parsimonious explanation is that ramping is not an intrinsic property of the preparatory activity in ALM. Instead, ramping may be explained by an external input, which shifts the location of the discrete attractors (Fig. 7e, f, Extended Data Fig. 11a-i).

## Discussion

We performed a series of experiments to probe the mechanisms underlying persistent preparatory activity in ALM. First, membrane potential dynamics and modulation of membrane potential were inconsistent with cell-autonomous mechanisms as a primary mechanism for persistent activity. Second, during the delay epoch, activity funneled toward two discrete endpoints, both at the level of membrane potential and spike rate, consistent with discrete attractor dynamics, but not with integrators. Third, after perturbations, detailed activity trajectories recovered to reach one of the two discrete endpoints, again consistent with discrete attractor dynamics. Fourth, when delay duration was randomly varied, activity during the delay epoch was approximately stationary, showing that ramping is not a necessary component of preparatory activity. These experiments provide direct evidence for discrete attractor dynamics as a mechanism underlying short-term memory.

How does ALM form a network with discrete attractors dynamics? ALM neurons with the same selectivity show high spike count correlations ^16^, implying preferential coupling among neurons sharing the same selectivity ^59^. Together with synaptic or postsynaptic non-linearities, such a network is expected to create discrete attractors without fine-tuning ^21^. In addition to the local excitatory connections, ALM is bidirectionally connected with several thalamic nuclei that show similar preparatory activity ^47^. Given the strong coupling between ALM and thalamus ^47^, our bilateral perturbations of ALM likely modified activity not only in ALM but also in these thalamic nuclei. The most parsimonious explanation is that attractor dynamics is generated by ALM within a cortico-thalamocortical loop.

One challenge in reconciling the measured activity with attractor models are large and systematic changes in neuronal activity during the delay epoch observed in this and other studies ^32,34^, whereas standard attractor models generate stationary activity once the network converged to the attractor. The disappearance of ramping up in the random delay task (Fig. 7) indicates that complex dynamics during the delay is not a necessary component of preparatory activity ^57^.

An additional question is how models should incorporate dynamics such as ramping ^32,34^. Ramping activity observed in the fixed delay task could be due to a non-selective external input to the network, reflecting for instance an urgency or timing signal ^57^ (Extended Data Fig 11b-i). Alternatively, ramping could be due to intrinsic aspects of the network dynamics, for instance a slow convergence towards the attractor (Extended Data Fig 11j-q). The coexistence of a slow ramping dynamics during the delay with a relatively fast recovery from perturbation, and the absence of ramping in some of our experiments, suggest that ramping is inherited rather than internally generated. Additional experiments are needed to clearly identify the source and the role of the ramping dynamics.

In contrast to transient bilateral perturbations of ALM, transient unilateral perturbation early in the delay epoch has no behavioral effect ^19^. Consistently, activity and selectivity recovers rapidly after unilateral perturbation. Recovery relies on input from the contralateral ALM via the corpus callosum ^19^. Previous studies have shown that multiple mechanisms, including integrators and discrete attractors, can be combined with a modular architecture to explain the robustness to unilateral inactivation. Models in which modular discrete attractor networks are distributed across the two hemispheres and coupled via the corpus callous account for all experimental results in this paper and also those in Li / Daie et al (2016) ^19^ (Extended Data Fig. 11).

Our task design only has two behavioral choices. This likely explains two stable endpoints in ALM dynamics. The attractor model can accommodate a large range of endpoints ^60^. It is possible that each learned movement corresponds to a discrete attractor. Testing this hypothesis represents an important area for future investigation.

Previous studies have reported evidence for discrete attractor dynamics. First, selective persistent activities in PFC of primates are robust to sensory distractors ^61,62^, and remain discrete even in response to continuously varying sensory stimuli ^63–65^. Our direct manipulation of ALM activity confirms that this robustness and discreteness are internal properties of circuit involving frontal cortex. Second, in a rat frontal cortex (FOF), behavioral effects of perturbation are consistent with discrete attractor dynamics ^66^. Third, across-trial variability of spike rate decays in a primate motor planning task ^52,53^. This led to the idea of “optimal initial conditions” required for specific movements ^15^. Our whole-cell recordings confirmed that such convergence of spike rates indeed reflects variance changes underlying membrane potential. Altogether, we propose that flexible yet robust discrete attractor dynamics subserves short-term memory in frontal cortex in a wide-range of behaviors.

## Acknowledgements

We thank Drs. N. Brunel, S. Lim, N. Li, M. Economo, K. Daie, J. Yu, T. Wang, L. Liu, A. Ebihara, and A. Finkelstein for comments on the manuscript, M. Inagaki for animal training, T. Harris, B. Barbarits, J. J. James and W. L. Sun for help with silicon probe recordings and spike sorting, and D. Hansel and S. Druckmann for discussions. This work was funded by Howard Hughes Medical Institute. H.K.I is a Helen Hay Whitney Foundation postdoctoral fellow.

## Author Contributions

H. K.I. performed experiments and analyzed data, with input from F.L., S.R. and K.S. F.L. and S.R. performed network modeling. H.K.I. and K.S. wrote the paper, with input from all the authors.

## Author Information

The authors declare no competing interests. Correspondence and requests for materials should be addressed to.

## Extended Data Figures Legends

**Extended Data Figure 1.**
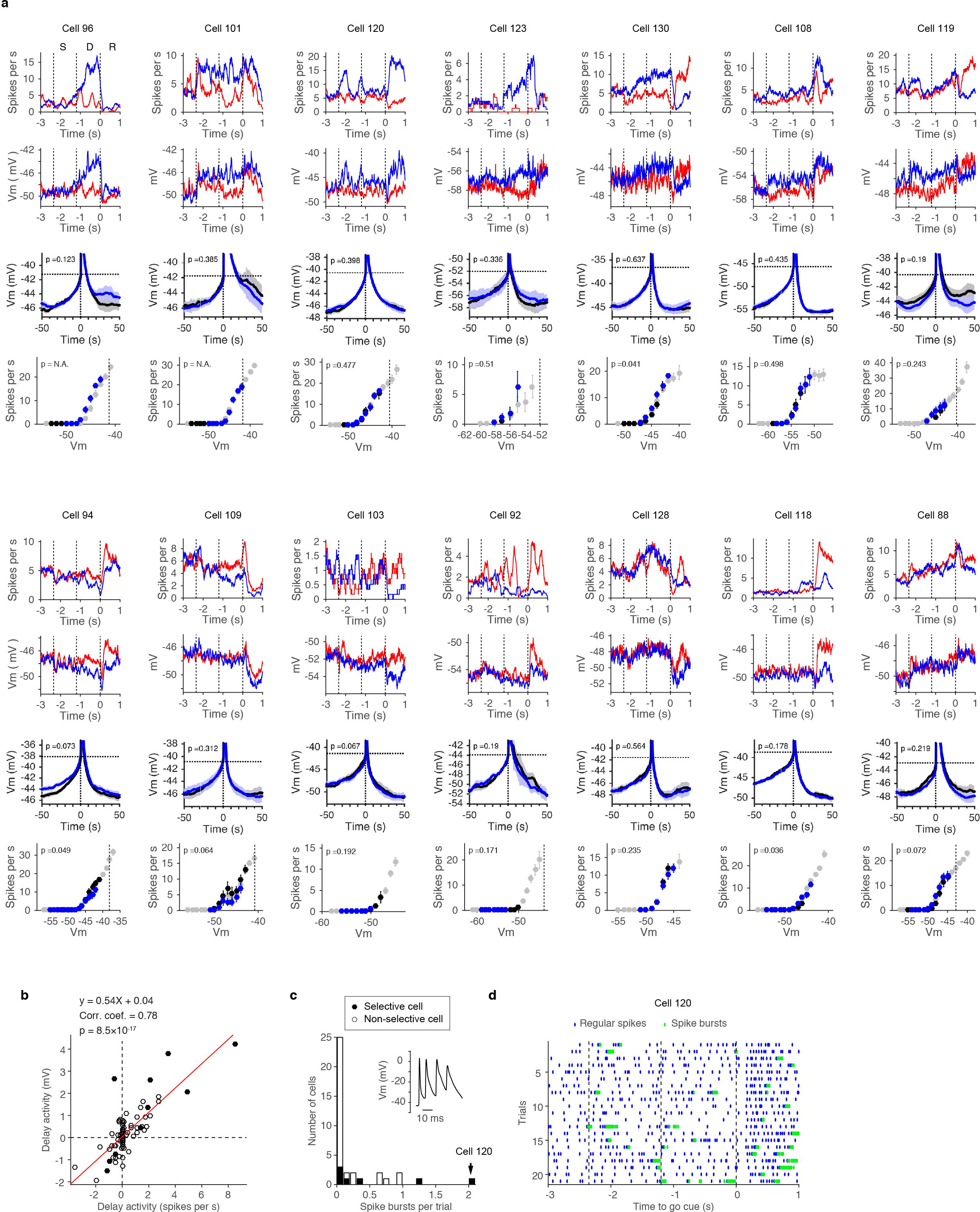
Whole-cell recordings during the auditory task. **a.** Example cells. Each column represents data from each cell. Top, mean SR; 2^nd^ row, mean Vm; 3^rd^ row, spike-triggered median of Vm. Spikes in the pre-sample epoch (black) and the delay epoch (blue) in the contra trials were analyzed. Shadow, S.E.M. (*hierarchical bootstrap*). *P*-value, probability of null hypothesis that spikes shapes are the same between epochs (Methods, *hierarchical bootstrap);* 4^th^ row, relationship between Vm and SR (Vm-to-SR curve). Vm-to-SR curves of the pre-sample epoch (black), the delay epoch (blue), and all epochs (grey) are shown. Bar, S.E.M. (*hierarchical bootstrap*). *P*-value, probability of null-hypothesis that Vm-to-SR curves are the same between the presample and delay epoch (Methods, *hierarchical bootstrap*). When there was no overlap between the curves from the two epochs, we did not test (*p* = N.A). **b.** Difference in SR and Vm between the pre-sample epoch and the delay epoch (“Delay activity”). For each cell, contra and ipsi trials are shown separately (n = 37). Black, selective cells (*n* = 10). Red line, linear regression. Linear regression, *Pearson* correlation coefficient (Corr. coef.), and the *t-statistic* of *Pearson* correlation coefficient (*p*) are shown. **c.** Distribution of number of spike bursts per trial. White, all cells; black, selective cells. Inset, an example spike bursts. **d.** Spike raster of contra trials in an example cell with high occurrence of spike bursts (cell 120, arrow in c). Blue, regular spikes; green, spikes belonging to spike bursts.

**Extended Data Figure 2.**
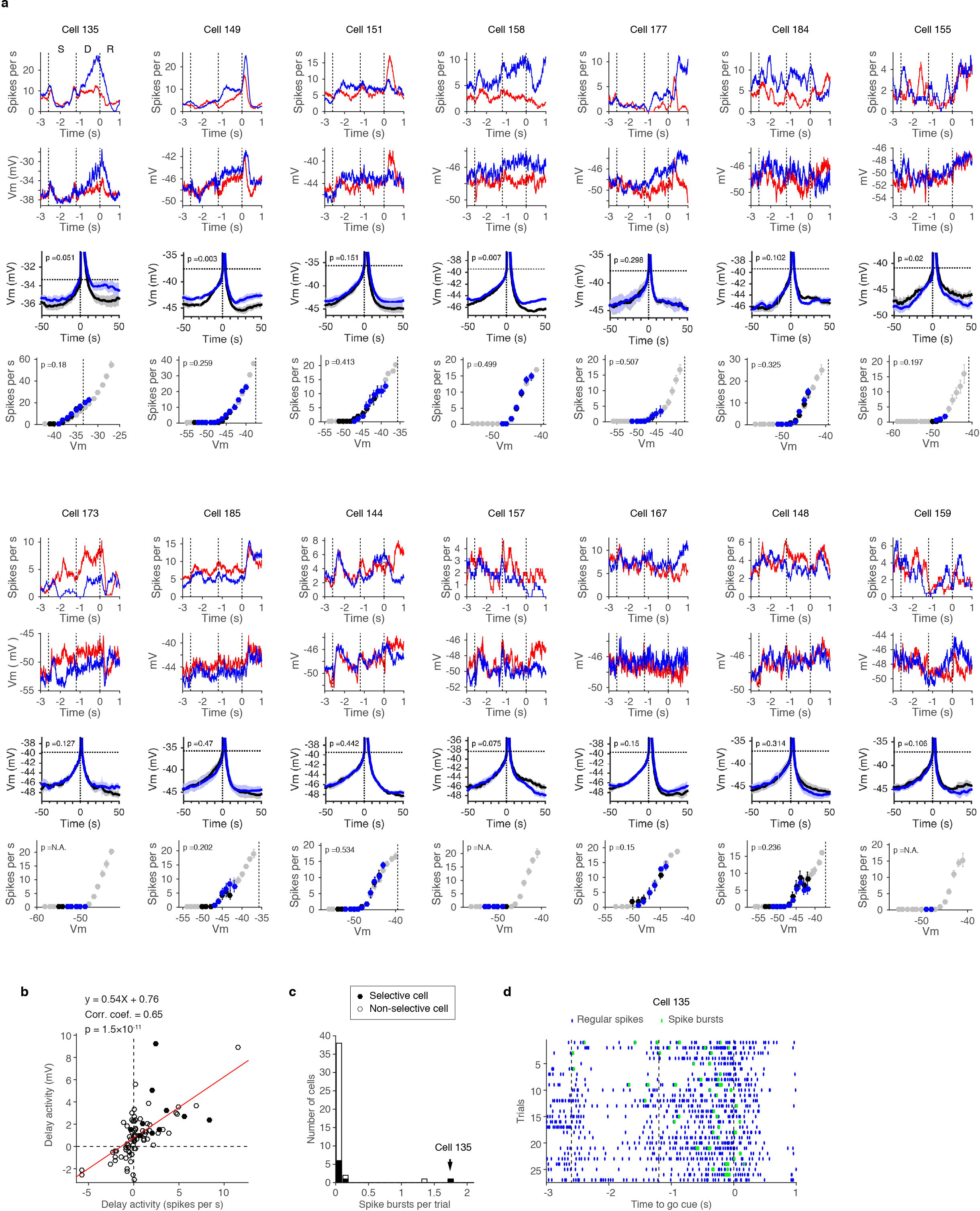
Whole-cell recordings during the tactile task. **a-d.** The same format as in **Extended Data Fig. 1** for cells recorded during the tactile task. All cells, *n* = 42; selective cells, *n* = 10.

**Extended Data Figure 3.**
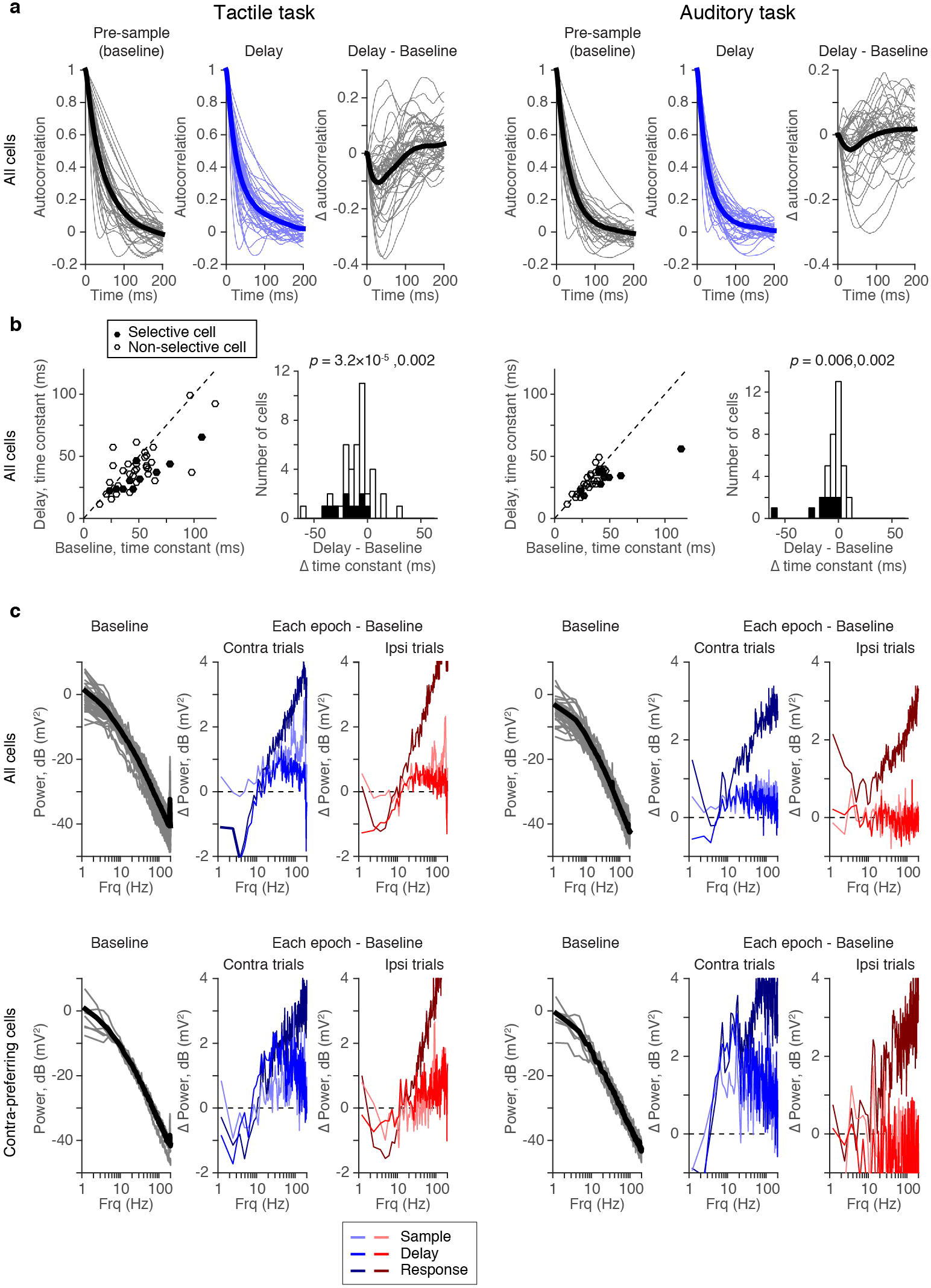
Autocorrelation and power spectral density of Vm. **a.** Autocorrelation of Vm during the pre-sample (baseline) epoch (left) and the delay epoch in contra trials (middle). Subtraction of these two autocorrelation curves are shown on right to emphasize the difference between epochs. Thick line, mean across cells. Thin lines, individual cells. Shorter time constant during the delay epoch is presumably due to increase in conductance. **b.** Comparison of time-constant of membrane fluctuations based on autocorrelation between the pre-sample epoch and the delay epoch. Left, scatter plot of the time constant; Right, histogram of the difference in time-constant between the delay epoch compared to the pre-sample epoch (Δ time constant). *P*-value, *signrank* test examining a null hypothesis that Δ time constant is 0. The first *p*-value, all cells (left); second *p*-value, selective cells (right). **c.**Power spectral density of Vm. Left, Power spectral density of Vm during the pre-sample epoch; Middle and right, Change in spectrum density in each epoch compared to the presample epoch. Different colors indicate different epochs (box).

**Extended Data Figure 4.**
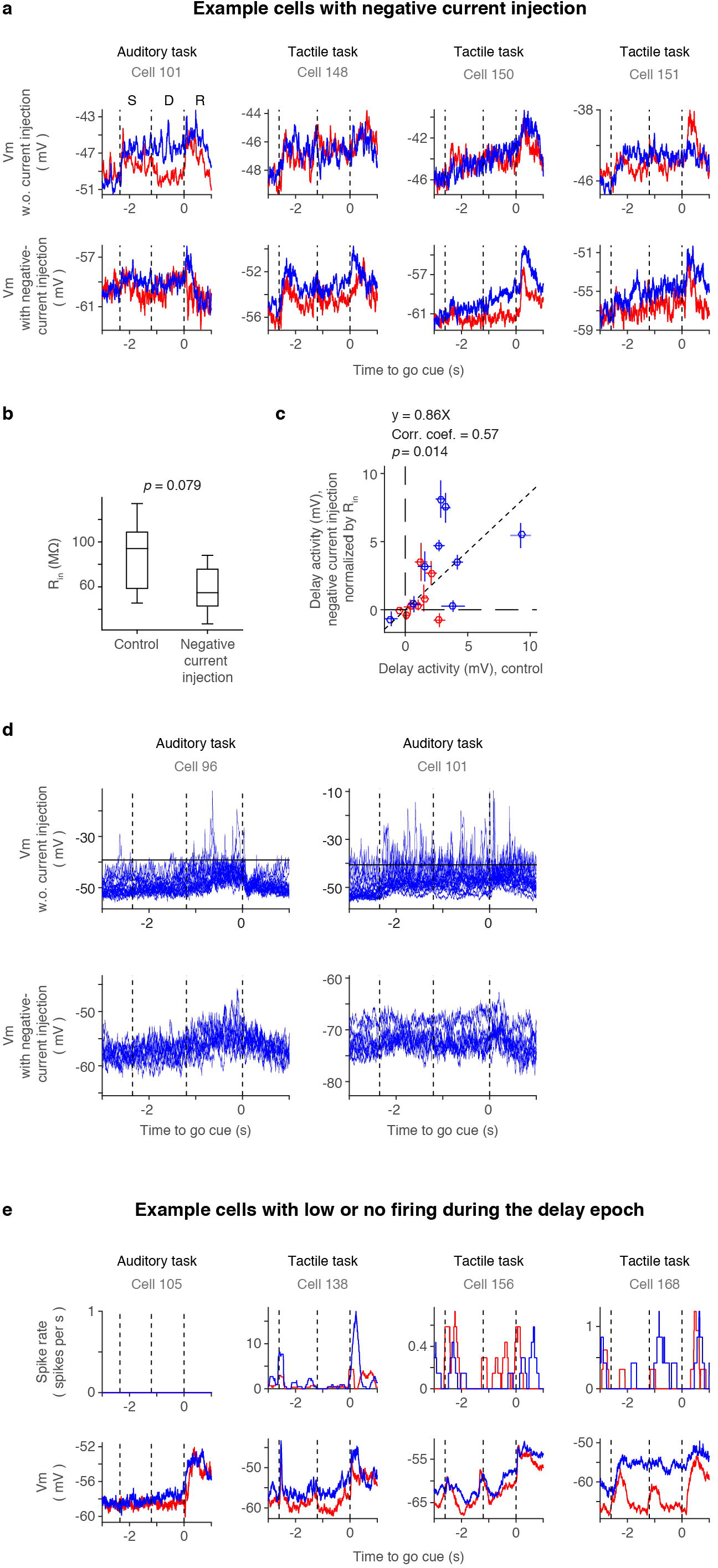
Negative current injection. **a.** Example cells with negative current injection. Top, mean Vm without negative current injection; Bottom, mean Vm with negative current injection. **b.** Input resistance (R_in_) of cells with and without current injection (*n* = 10). Data was pooled from cells analyzed in **c**. **c.** Delay activity of Vm (increase in Vm from the pre-sample epoch to the delay epoch) with and without current injections (*n* = 10). Delay activity during the current injection was normalized by the change in input resistance. Contra trials (blue) and ipsi trials (red) are shown separately. Crosses, S.E.M. (*bootstrap*). Dashed line, linear regression. Slope of linear regression, *Pearson* correlation coefficient (Corr. Coef.), and the *t-statistic* of *Pearson* correlation coefficient (*p*) are shown. **d.** Loss of spike bursts after current injection. Top, Vm of two example cells with high occurrence of spike bursts. Overlay of all contra trials. Black horizontal line, spike threshold. Regular spikes were trimmed and Vm was averaged over 100 ms in this plot. Sharp overshoots above the spike threshold indicate spike bursts (Method). Bottom, Vm of the same example cells with negative current injection. There was no spike to trim. Vm was averaged over 100 ms. Note the loss of sharp depolarizing events. **e.** Example cells without spikes or with low SR during the delay epoch. Top, mean SR; 2^nd^ row, mean Vm. All of these cells showed significant difference in Vm between the contra and ipsi trials (*ranksum test*, *p* <0.05).

**Extended Data Figure 5.**
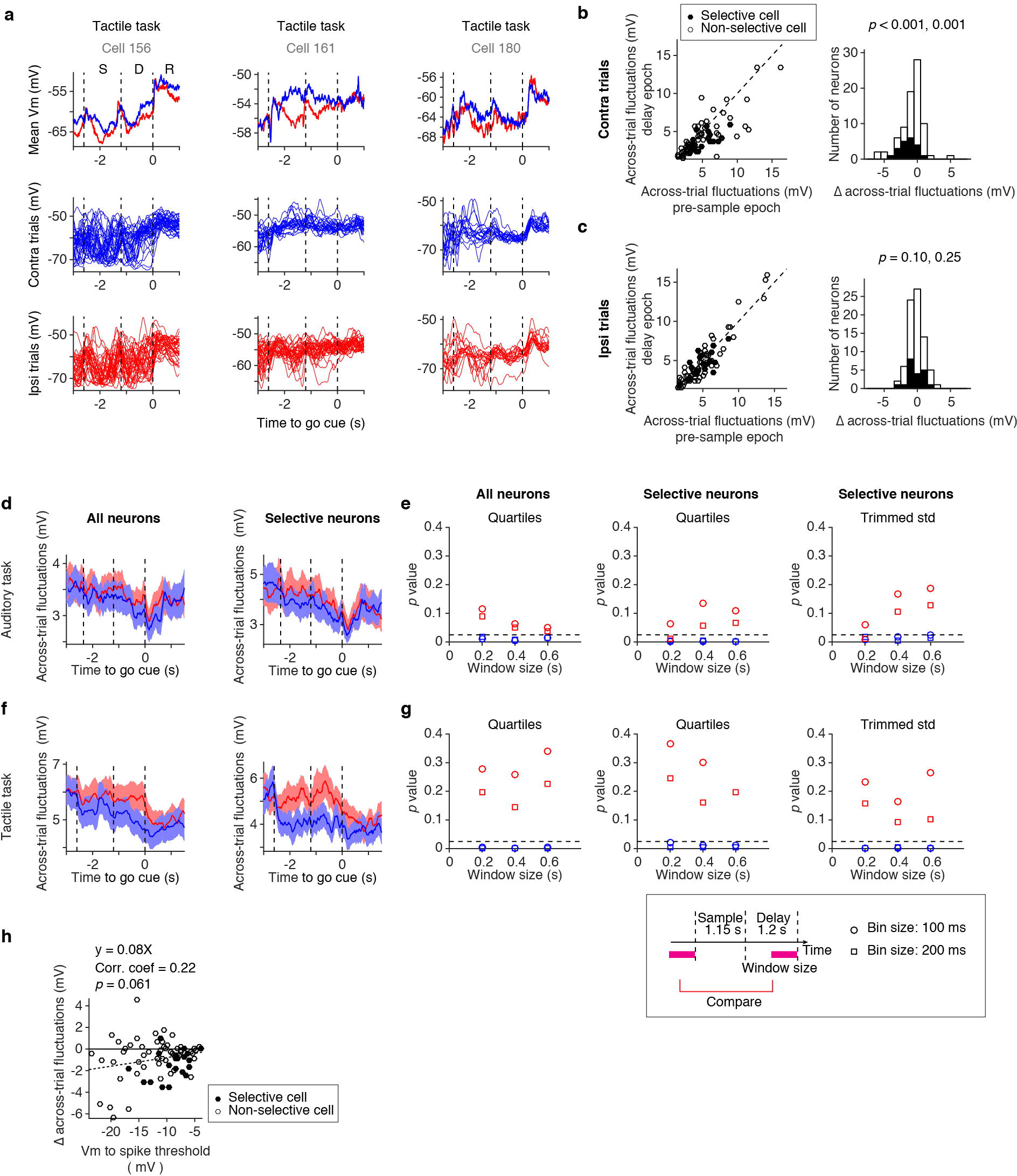
Funneling of Vm. **e.** Three example ALM neurons. Top, mean Vm; middle, all contra trials overlaid; bottom, all ipsi trials overlaid. To remove fast within trial fluctuations, Vm was averaged over 200 ms. **f.** The same plot as in **Fig. 4c, d**. **g.** Same as **b** for the ipsi trials. **h.** Across-trial fluctuations of all cells (*n* = 37) (left), and selective cells (*n* = 10) (right) in the auditory task (same as **Fig. 4b**). Line, mean of across-trial fluctuations; Shadow, S.E.M. (*hierarchical bootstrap*). **i.** Testing the decrease in across-trial fluctuations. *P*-value indicates the probability of a null-hypothesis that across-trial fluctuations during the delay epoch was higher than that during the pre-sample epoch (*hierarchical bootstrap*, *n* =1000 iteration). Across-trial fluctuations was measured as the quartile difference (left) or trimmed-standard deviation (right) of Vm across the same trial type (blue, contra trials; red, ipsi trials) (Methods). Both methods provided similar results. Vm was averaged over 100 (triangle), or 200 (square) ms to remove fast within trial fluctuations. The result was robust to the averaging bin size. The dashed line, *p* = 0.025 (*α* = 0.05 for two sided test). Box below **g**, Schematic of statistical test in **e** and **g**. Across-trial fluctuations during the pre-sample epoch (0.6 s) was compared with the across-trial fluctuations during the delay epoch (variable durations: window size in **e** and **g**). The window ends at the time *t*, which is *t* = (time of the go cue) - (bin size)/2, to exclude the signal after the go cue. **j.** Same as d for the tactile task. **k.** Same as e for the tactile task. **l.** Relationship between Vm and A across-trial fluctuations. Vm during the delay epoch was averaged and normalized to the spike threshold. Dashed line, linear regression. Slope of linear regression, *Pearson* correlation coefficient (Corr. Coef.), and the *t-statistic* of *Pearson* correlation coefficient (*p*) are shown. Pooling both tactile and auditory task (*n* =73). Note that the slope of regression line is opposite from what is expected for the ceiling effect.

**Extended Data Figure 6.**
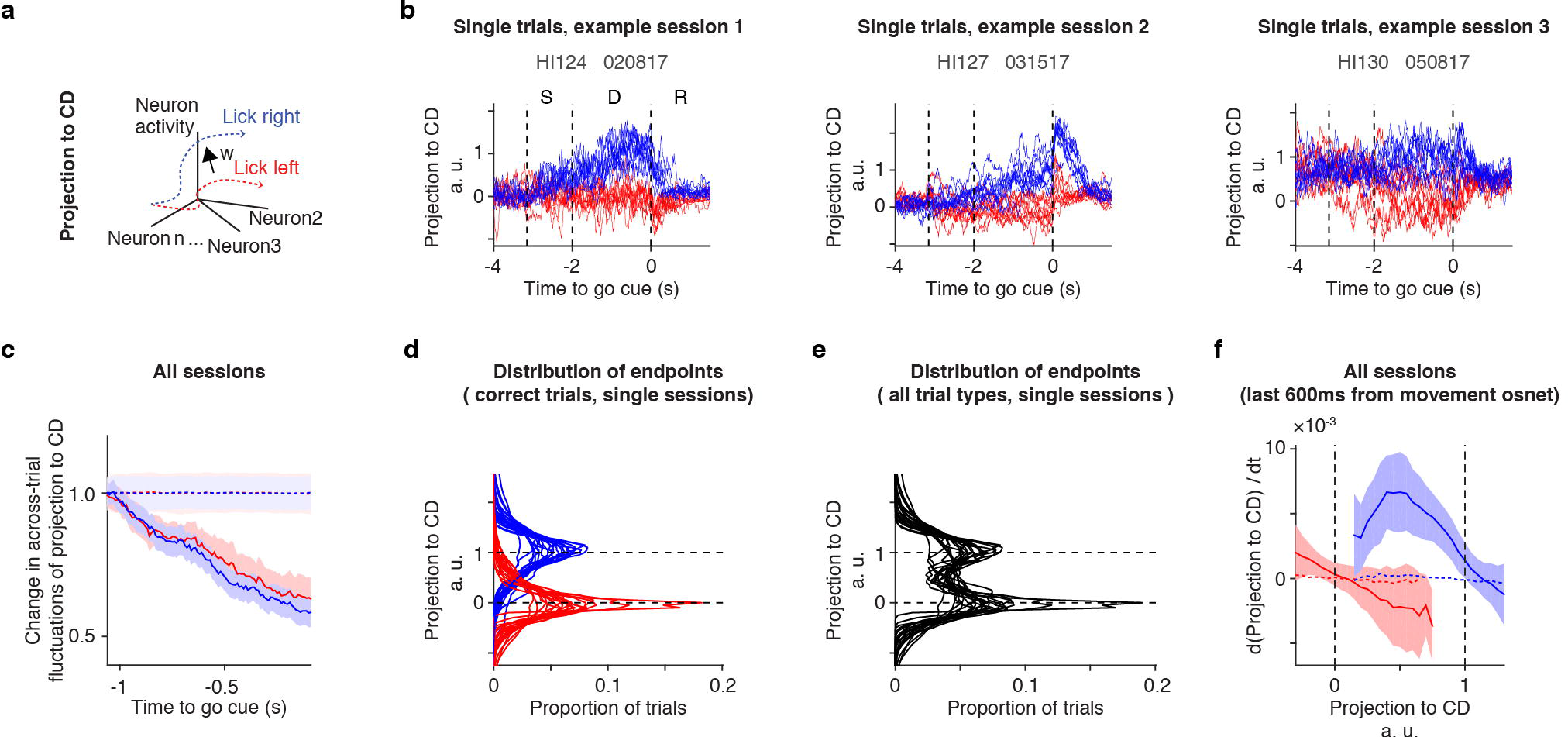
Characterization of funneling along CD. **a.** Schematics. Projection to coding direction (CD). **b.** Projection of trials to coding direction (CD) in example sessions. The same format as in **Fig. 5a**. **c.** Across-trial fluctuations in all sessions. Trials from all sessions were pooled. Solid line, grand mean. Dashed line, shuffled data (Methods). Shading, S.E.M. (*hierarchical bootstrap*). **d.** Distribution of endpoints (activity level at the last bin before go cue). Distribution of endpoints in each session is overlaid (*n* = 20 sessions). Distribution of contra trials (blue), ipsi trials (red) are shown. **e.** Distribution of endpoints of all trial types (both correct and incorrect trials, but not early lick or no-response trials). Distribution of endpoints in each session is overlaid (*n* = 20 sessions). **f.** Relationship between activity level along the CD, and the drift speed of trajectories. Solid line, data. Dashed line, shuffled data. Shading, S.E.M. (*hierarchical bootstrap*). Data of correct lick right trials (blue) and correct lick left trials (red) were analyzed separately. The same dataset as used in **Fig. 5f**.

**Extended Data Figure 7.**
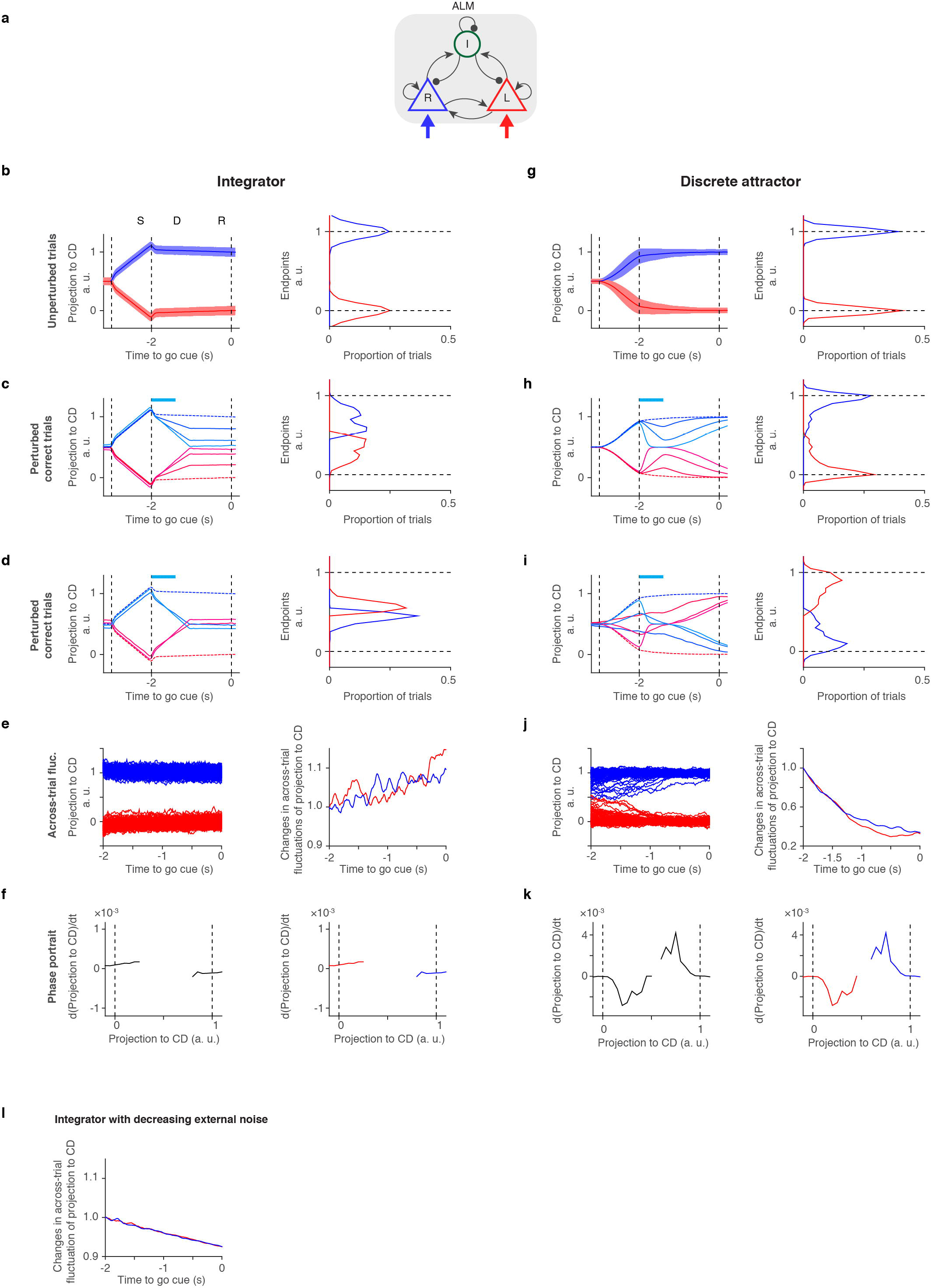
Discrete attractor and integrator models. **a.** Schematic of simulated one-hemisphere networks. **b-f**, integrator model. **g-k**, discrete attractor model. R, L, and I correspond to lick right selective excitatory neurons, lick left selective excitatory neurons, and inhibitory interneurons, respectively. Blue and red arrows, selective external input. **b.** Trajectories projected along CD. Activity in unperturbed trials (left), and distribution of endpoints (right). Line, mean; Shading, standard deviation. **c.** Activity in perturbed correct trials (left), and distribution of endpoints (right). Blue box on top, photoinhibition; Traces with lighter colors correspond to higher intensity of photoinhibition (Methods). **d.** Activity in perturbed incorrect trials (left), and corresponding distribution of endpoints (right). **e.** Trajectories during the delay epoch (1/6 of single trials are shown) (left), and across-trial fluctuations calculated based on these trajectories (right). Across-trial fluctuations are normalized for the value at −2.0 s before the go cue. **f.** Phase portrait of trajectories. Lick right and lick left trials were pooled together (left) or analyzed separately (right). **j-q.** Dynamics of one-hemisphere discrete attractor model. **l.** Across-trial fluctuations in integrator with decreasing external noise (Methods), normalized by their value 2.0 s before the go cue.

**Extended Data Figure 8.**
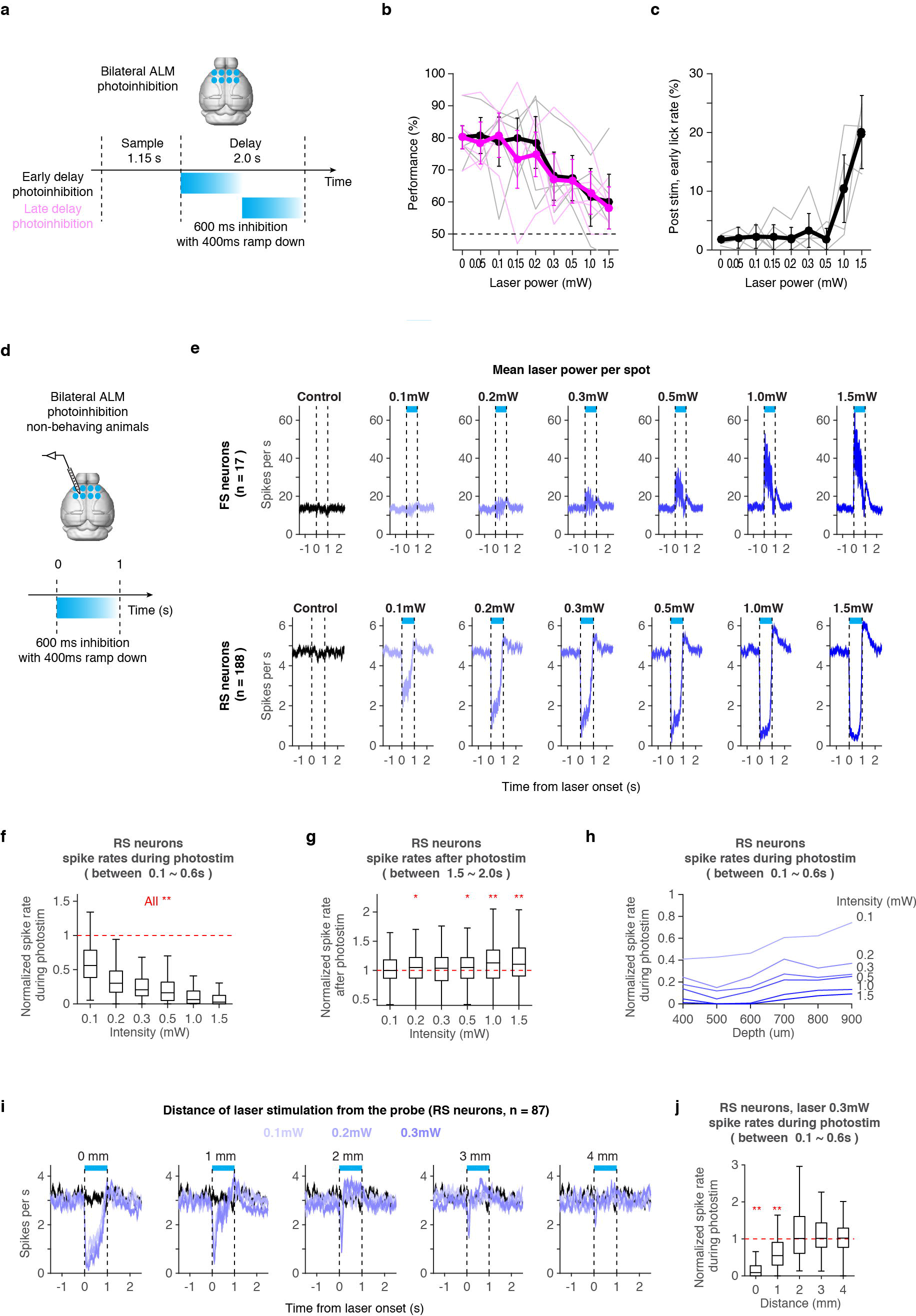
Characterization of bilateral photoinhibition of ALM. **a.** Schematic of bilateral photoinhibition of ALM. Eight spots surrounding ALM in both hemispheres (1 mm interval) were photoinhibited to silence whole ALM. Photoinhibition started at the onset of the delay epoch (−2 s to the go cue) or at the middle of the delay epoch (−1 s to the go cue) and lasted for 600 ms with 400 ms ramping down (Methods). **b.** Effect of the photoinhibition with different laser power on behavioral performance. Black, early delay inhibition; magenta, late delay inhibition. Thick lines, grand mean performance (*n* = 5 animals); thin lines, mean performance of each animal. Error bar, S.E.M (*hierarchical bootstrap*). Laser power is a mean power per spot. **c.** Effect of the photoinhibition with different laser power on early lick rates. Early lick rate after the early delay photoinhibition is shown. The same format as in **b**. **d.** Schematic of silicon probe recording during bilateral photoinhibition of ALM in non-behaving animals (relevant to **e-j**). **e.** SR of fast spiking (FS) neurons (top) and regular spiking (RS) neurons (bottom) during the eight spots photoinhibition of ALM. Each column represents data with different laser power. Mean SR is shown. Blue box, photoinhibition. **f.** SR of RS neurons during the photoinhibition. Mean SR during the photoinhibiton (100 to 600 ms) was divided by mean SR before the photoinhibiton (−1 to 0 s) to calculate the normalized SR in each cell (Same for **h** and **j**). Increasing laser power resulted in stronger inhibition. **: *p* < 0.01, *signrank test* with *Bonferroni* correction. From left to right *p* = 2.1×10^−19^, 2.2×10^−25^, 2.1×10^−27^, 4.2×10^−27^, 1.6×10^−27^, 2.7×10^−31^ (*p*-values without *Bonferroni* correction), *n* = 188. **g.** SR of RS neurons after the photoinhibition. Mean SR after photoinhibiton (1 to 2 s) was divided by the mean SR before the photoinhibiton (−1 to 0 s) to calculate normalized SR in each cell. Increasing laser power resulted in stronger rebound. *: *p* < 0.05, **: *p* < 0. 01, *signrank test* with *Bonferroni* correction. From left to right *p* = 0.37, 7.1×10^−3^, 3.1×10^−2^, 1.4×10^−2^, 3.6×10^−5^, 1.4×10^−6^ (*p*-values without *Bonferroni* correction), *n* = 188. **h.** Relationship between depth and SR of RS neurons during the photoinhibition. SR were averaged based on the depth of neurons. Photoinhibition affected neurons across layers. **i.** SR of RS neurons during the photoinhibition at different locations. Mean SR is shown for each laser power (control, black). Distance of recording site (ALM) and center of laser stimulation (center of four spots in the same hemisphere) is shown. Laser spots were moved from ALM to posterior locations. **j.** SR of RS neurons during the photoinhibition at different locations. Photoinhibition affected neurons 1 mm away from the laser, consistent with previous report ^11^. ** *p* < 0.01, *signrank test* with *Bonferroni* correction. From left to right *p* = 1.1×10^−15^, 6.0×10^−7^, 0. 61, 0.55, 0.52 (*p*-values without *Bonferroni* correction), *n* = 87.

**Extended Data Figure 9.**
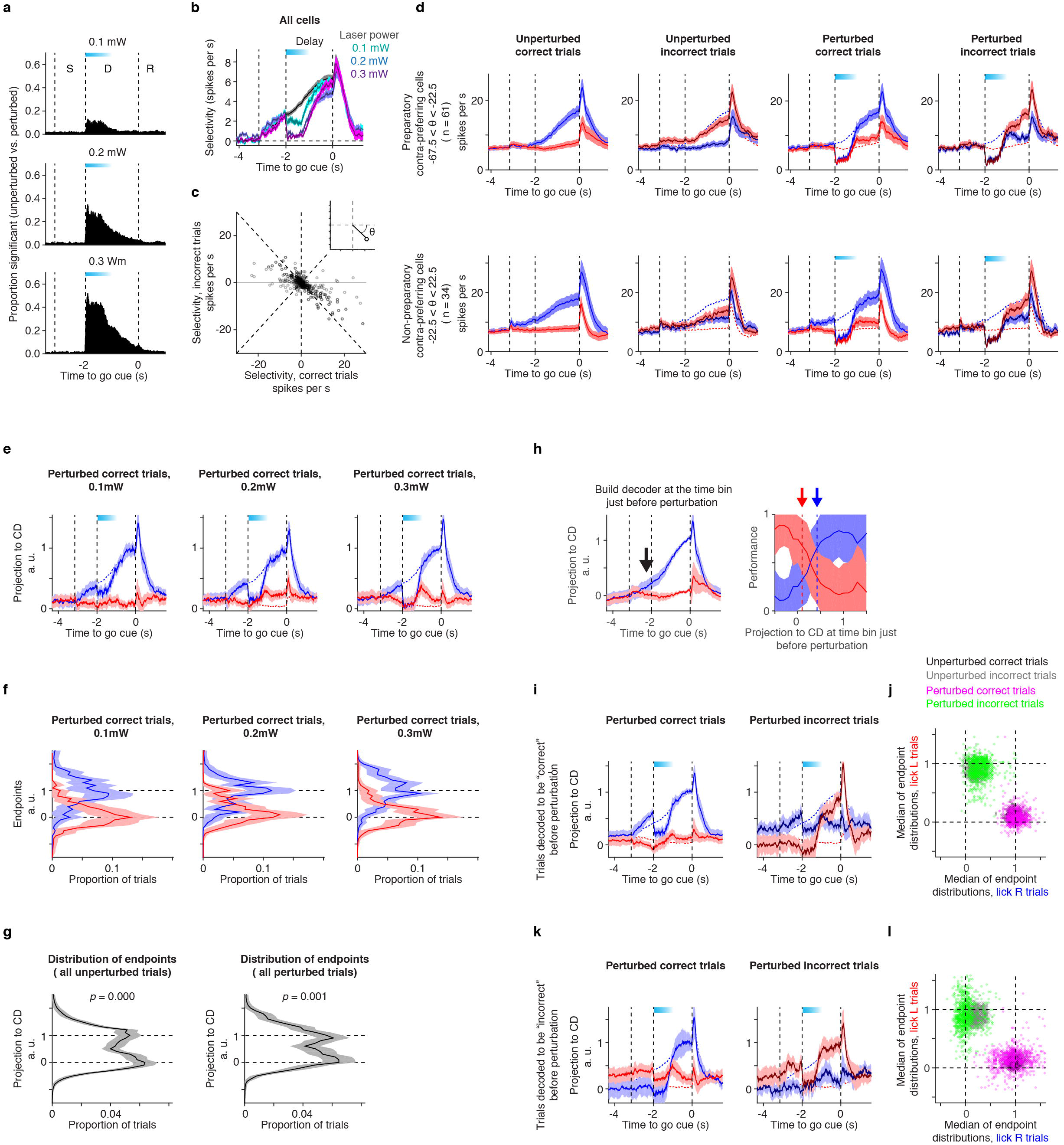
Robustness of trajectories to the photoinhibition. **a.** Proportion of cells with significant SR difference between unperturbed and perturbed trials at each time point (Methods). Blue box, photoinhibition. **b.** Selectivity during bilateral photoinhibition. Line, mean of all cells; Shadow. S.E.M (*bootstrap*). Black, unperturbed trials. Blue box, photoinhibition. **c.** Relationship between selectivity in correct and incorrect trials. Mean selectivity during the last 600 ms of the delay epoch is shown. Inset, definition of *θ* (Methods). **d.** Mean SR of contra-preferring neurons with different *θ*. Lines, grand mean of peri-stimulus time histogram (PSTH); Shading, S.E.M. (*bootstrap*). Mean PSTHs of unperturbed correct trials (1^st^ row) are also shown as dashed lines in 2^nd^ to 4^th^ row. **e.** Trajectories along CD for correct trials photoinhbitied with different laser intensities. The same format as in **Fig. 6e**. Trials from all sessions were pooled. **f.** Distribution of endpoints in e. Shading, S.E.M. (*hierarchical bootstrap*). Distributions of contra trials (blue) and ipsi trials (red) are shown. **g.** Distribution of endpoints in Fig. 6e pooling all trial types (both correct and incorrect trials, but not early lick or no-response trials). Unperturbed (left) and perturbed (right) trials. Shading, S.E.M. (bootstrap). P-value indicates the chance that proportion of trials at the middle of two endpoints (Projection to CD = 0.5) are higher than the mean proportion of trials at the endpoints (projection to CD = 0 and 1) (*hierarchical bootstrap*, *n* = 1000 iteration). Low *p* value indicates bimodal distribution. **h.** Decoding of licking direction based on projection to CD just before perturbation. Left, we developed a decoder based on unperturbed trials (Methods). Arrow, the time bin used for decoding (the last bin before perturbation); Right, relationship between projection to CD and performance in lick right (blue) or left (red) trials. Dashed lines and arrows indicate decoders (the point crossing 70 % performance) for contra (blue) and ipsi (red) trials. Shading, S.E.M. (bootstrap). **i.** Trajectories along CD in trials decoded to be “correct trials” before the onset of perturbation. The same format as in **Fig. 6e**. **j.** Median of the endpoint distributions in **i**. The same format as in **Fig. 6h**. **k.** Trajectories along CD in trials decoded to be “incorrect trials” before the onset of perturbation. The same format as in **Fig. 6e**. **l.** Median of the endpoint distributions in **k**. The same format as in **Fig. 6h**.

**Extended Data Figure 10.**
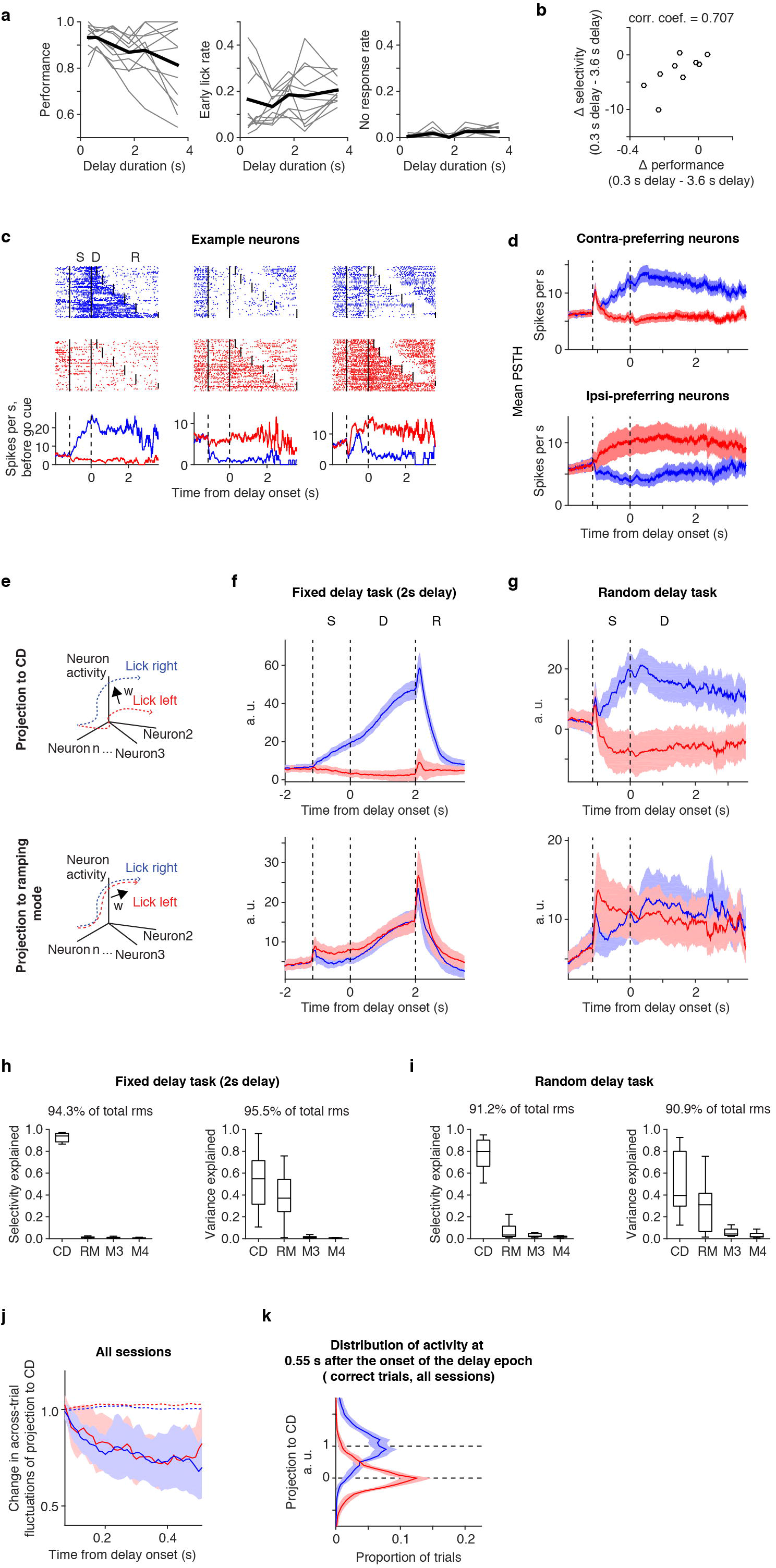
Performance in the random delay task. **a.** Behavioral performance (left), early lick rate (middle) and no-response rate (right) in the random delay task. Thin lines, individual sessions (*n* = 11). Thick line, grand mean among sessions. **b.**Relationship between decay in performance (Δ performance) and decay in selectivity (Δ selectivity) during the delay epoch. Both decays were calculated as differences between trials with the shortest delay (0.3 s) and the longest delay (3.6 s). Individual circle represents individual session. Corr. Coeff., *Pearson* correlation coefficient. Selectivity during the delay epoch had a high correlation with performance, which indicates that the decay in selectivity may reflect loss of short-term memory over time. **c.** Example cells. Top, spike raster. Dashed lines separate epochs. Ten trials per delay duration were randomly selected; Bottom, mean SR based on spikes before the go cue across trials with different delay durations. Because delay durations are different among trials, time axis is aligned to the onset of the delay epoch (same in **d, g**). **d.** Grand mean SR of contra-preferring neurons (top) and ipsi-preferring neurons (bottom). Shadow, S.E.M. (*bootstrap*). **e.**Schematics. Projection of trials to CD (top) and ramping mode (RM) (bottom). **f.**Projection of trials to CD (top) and RM (bottom) in fixed delay task with 2 s delay. Time axis is aligned to the onset of the delay epoch to be consistent with **g**. Line, grand mean across sessions. S.E.M. *bootsrap* across sessions. CD is not normalized to the endpoints (unlike in Fig.6). **g.** The same as in f for the random delay task. **h.** Selectivity during the delay epoch explained by each mode (left), and variance during the delay epoch explained by each mode (right) in fixed delay task with 2 s delay. RM, ramping mode. M3 and M4, third and fourth SVD component (Methods). Sum of four modes are shown on top (mean across sessions). **i.** The same as in **h** for the random delay task. **j.** Across-trial fluctuations in all sessions. Trials from all sessions were pooled (*hierarchical bootstrap*). The same format as in Extended Data Fig. 6c. **k.** Distribution of activity at 0.55 s after the onset of the delay epoch (we did not include trials with 0.3 s delay). The same format as in Fig. 5c.

**Extended Data Figure 11.**
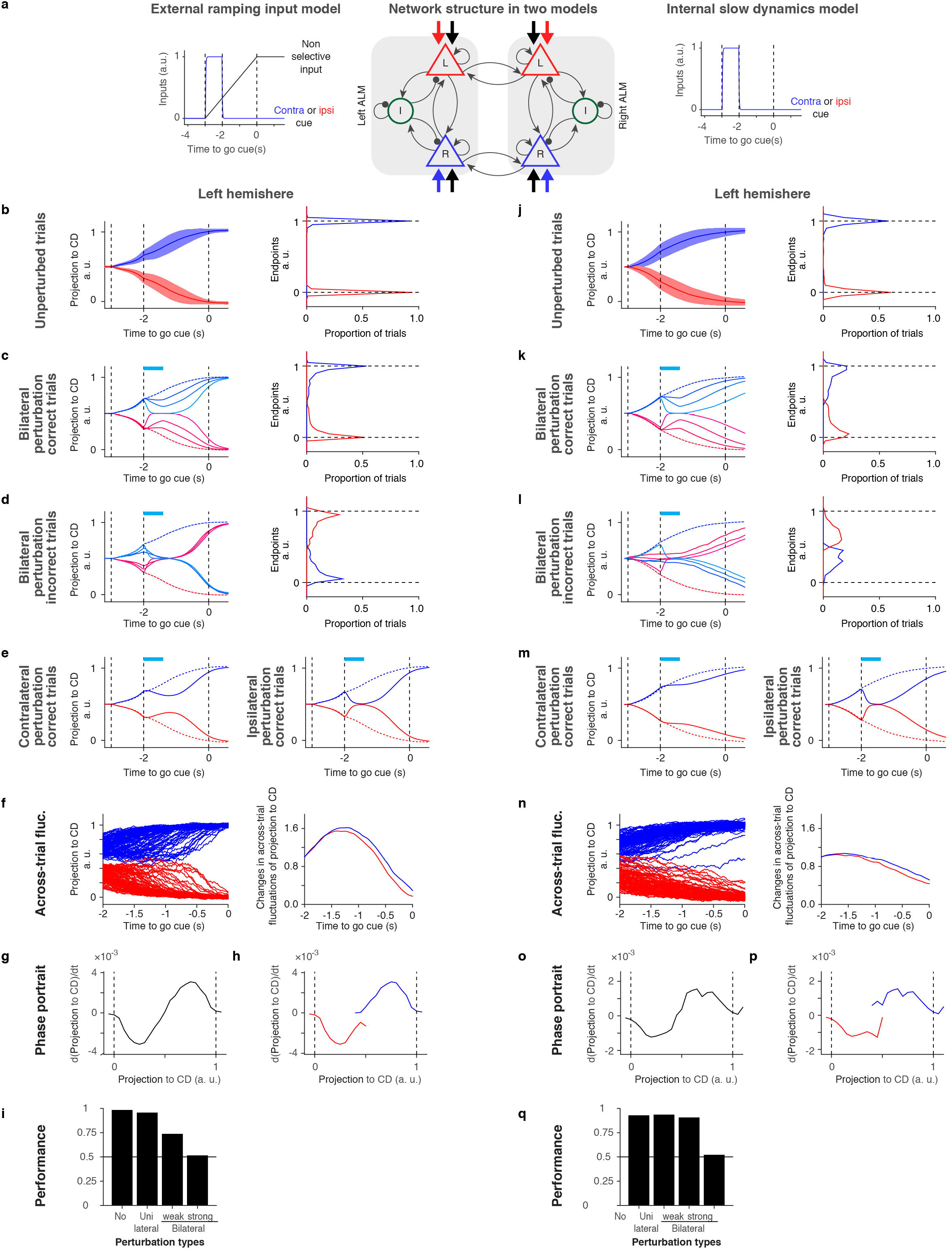
Discrete attractor models with two hemispheres. We built two-hemisphere models in order to explain 1) ramping dynamics during the delay epoch, and 2) effect of unilateral ALM perturbation described in Li et al ^19^. The circuit architecture in each hemisphere is the same as in Extended Data Fig.7. Slow ramping was caused by external non-selective input (**b-i**) or internal slow dynamics (**j-q**). All panels contain results from the left hemisphere. **a.** Schematic of two-hemisphere network models. Circuit architecture, common to both models (middle), and time course of external inputs in each model (Left and Right). R, L, and I correspond to lick right selective excitatory neurons, lick left selective pyramidal neurons, and inhibitory interneurons, respectively. Blue and red arrows, selective external input. Black arrows, non-selective external input. **b.** Trajectories projected along CD in model with external ramping input. Activity in unperturbed trials (left), and distribution of endpoints (right). Line, mean; shading, standard deviation. **c.** Projected activity in bilateral photoinhibition correct trials (left), and distribution of endpoints (right). Line, mean. Blue box on top, photoinhibition. As in Extended Data Fig, 7, lighter colors correspond to higher intensity of photoinhibition (Methods). **d.** Projected activity during bilateral photoinhibition incorrect trials. **e.** Projected activity during unilateral photoinhibition correct trials. Effect of unilateral ALM photoinhibiton contralateral (left panel) or ipsilateral (right panel) to the analyzed hemisphere (left ALM) are shown. **f.** Trajectories during the delay epoch (1/6 of all trials are shown) (Left), and across-trial fluctuations during the delay epoch (Right), normalized by their value 2.0 s before the go cue. **g.** Phase portrait of projected trajectories. Contra and ipsi trials were pooled together. **h.** Phase portrait of trajectories. Contra and ipsi trials were analyzed separately. **i.** Behavioral performance of the model (methods). **j-q.** Same as in **b-i** for internal slow dynamics model.

## Methods

### Mice

This study is based on data from 26 male mice (age > P60). We used four transgenic mouse lines: PV-IRES-Cre ^67^, Ai32 (Rosa-CAG-LSL-ChR2(H134R)-EYFP-WPRE, JAX 012569) ^68^, Gad2-cre (a gift from Boris Zemelman), and VGAT-ChR2-EYFP ^69^ (See Supplementary Information Table 1-3 for detail).

All procedures were in accordance with protocols approved by the Janelia Institutional Animal Care and Use Committee. Detailed information on water restriction, surgical procedures and behavior have been published ^70^. Surgical procedures were carried out aseptically under 1-2 % isofluorane anesthesia. Buprenorphine HCl (0.1 mg/kg, intraperitoneal injection; Bedford Laboratories) was used for postoperative analgesia. Ketoprofen (5 mg/kg, subcutaneous injection; Fort Dodge Animal Health) was used at the time of surgery and postoperatively to reduce inflammation. After the surgery, mice were allowed free access to water for at least three days before start of water restriction. Mice were housed in a 12:12 reverse light:dark cycle and behaviorally tested during the dark phase. A typical behavioral session lasted 1 to 2 hours and mice obtained all of their water in the behavior apparatus (approximately 1 ml per day; 0.3 ml was supplemented if mice drank less than 0.5 ml). On other days mice received 1 ml water per day. Mice were implanted with a titanium headpost. For ALM photoinhibition, mice were implanted with a clear skull cap ^11^. Craniotomies for recording were made after behavioral training.

### Behavior

Mice were trained using established procedures ^70^. For the tactile task (Fig. 2-4), at the beginning of each trial, a metal pole (diameter, 0.9 mm) moved within reach of the whiskers (0.2 s travel time) for 1.0 second, after which it was retracted (0.2 s retraction time). The sample epoch (1.4 s total) was the time from onset of the pole movement to completion of the pole retraction. The delay epoch lasted for another 1.2 seconds after completion of pole retraction. An auditory ‘go’ cue separated the delay and the response epochs (pure tone, 3.4 kHz, 0.1 s).

For the auditory task (Fig. 2-7), at the beginning of each trial, five tones were presented at one of two frequencies: 3 or 12 kHz. Each tone was played for 150 ms with 100 ms intertone intervals. The sample epoch (1.15 s total) was the time from onset of the first tone to the end of the last tone. The following delay epoch lasted for another 1.2 (Fig. 2-4) or 2.0 (Fig.5-6) seconds. An auditory ‘go’ cue (carrier frequency 6 kHz, with 360 Hz modulating frequency, to make it distinct from instruction tones) separated the delay and the response epochs (0.1 s). To compensate the sound intensity for tuning curve of C57BL6 auditory system ^71^, the sound pressure was 80, 70, and 60 dB for 3, 6 and 12 kHz sound, respectively. These frequencies are relatively invulnerable to hearing loss observed in C57BL6 mice ^72^.

For the “random delay task” (Fig. 7), delay durations were randomly selected from six values (0.3, 0.6, 1.2, 1.8, 2.4, 3.6 s). Probability of the delay durations mimicked cumulative distribution function of the exponential distribution (**τ** = 0.9 s) with 0.3 s offset (mean delay duration: 1.2 s) (Fig. 7a). Because the hazard function of the exponential distribution is constant, animals cannot predict the timing of the go cue (or reward). Performance in this “random delay task” was similar to that in a task with a fixed delay duration, or “fixed delay task” (Extended Data Fig. 9a).

The “pre-sample” (baseline) epoch was 1.2 seconds long, unless otherwise described. A two-spout lickport (4.5 mm between spouts) was used to record licking events and deliver water rewards. After the ‘go’ cue, licking the correct lickport produced a water reward (approximately 2 (μL); licking the incorrect lickport triggered a timeout (0 to 5 s). Licking early during the trial (‘early lick’ trials) was punished by a timeout (1 s). Trials in which mice did not lick within 1.5 seconds after the ‘go’ cue (‘no response’ trials) were rare and typically occurred at the end of behavioral sessions. These ‘no response’ trials and ‘early lick’ trials were excluded from analyses. For the random delay tasks, mice were first trained with a fixed delay duration of 1.2 s. After criterion performance was reached (80 % correct), we switched to the random delay task and trained at least one additional week before recordings.

### Photoinhibition

Photoinhibition was deployed on 25-33 % behavioral trials (Fig. 6 and Extended Data Fig. 9). To prevent mice from distinguishing photoinhibition trials from control trials using visual cues, a ‘masking flash’ (1 ms pulses at 10 Hz) was delivered using 470 nm LEDs (Luxeon Star) throughout the trial. Photostimuli from a 473 nm laser (Laser Quantum) were controlled by an acousto-optical modulator (AOM; Quanta Tech) and a shutter (Vincent Associates).

Photoinhibition of ALM was performed through the clear-skull cap (beam diameter at the skull: 400 μm at 4 σ) ^11^. The light transmission through the intact skull is 50 %. We stimulated parvalbumin-positive interneurons in PV-IRES-Cre mice crossed to Ai32 reporter mice expressing ChR2. Behavioral and electrophysiological experiments showed that photoinihibition in the PV-IRES-Cre x Ai32 mice was indistinguishable from the VGAT-ChR2-EYFP mice ^47^.

To silence ALM bilaterally during early or late delay (Fig. 6 and Extended Data Fig. 9), we photostimulated for 0.6 s (40 Hz photostimulation with a sinusoidal temporal profile) with 0.4 s ramping down, starting at the beginning of the delay epoch or 1.0 s after the beginning of the delay epoch, respectively. We photostimulatd four spots in each hemisphere, centered on ALM (AP 2.5 mm; ML 1.5 mm) with 1 mm spacing (in total eight spots bilaterally) using scanning Galvo mirrors. We photoinhibited each spot sequentially at the rate of 5 ms per step. The laser power noted in the figures and text indicates the mean laser power per spot. The total laser power was therefore eight-fold higher. The bilateral manipulation prevents rescue of neural dynamics from unaffected regions ^19^.

### Behavioral data analysis

Behavioral performance was the proportion of correct trials, excluding ‘lick early’ and ‘no response’ trials (Extended Data Fig. 8b and 10a). Behavioral effects of photoinhibition were quantified by comparing the performance with photoinhibition with control performance. Early lick rate (Extended Data Fig. 8c and 10a) was the proportion of early lick trials excluding no response trials. For Extended Data Fig. 8c, only the early licks during the last 1 s of the delay epoch (post-photoinhibition) were counted. No response rate (Extended Data Fig. 10a) was the proportion of no response trial.

### In vivo whole-cell recording

All recordings were made from the left hemisphere. Whole-cell recordings were made using pulled borosilicate glass (Sutter instrument, CA). A small craniotomy (100 - 300 μm diameter) was created over ALM under isofluorane anesthesia and covered with cortex buffer (125 mM NaCl, 5 mM KCl, 10 mM glucose, 10 mM HEPES, 2 mM MgSO4, 2 mM CaCl2. Adjust pH to 7.4). Whole-cell patch pipettes (7-9 MΩ) were filled with internal solution (in mM): 135 K-gluconate, 4 KCl, 10 HEPES, 0.5 EGTA, 10 Na_2_-phosphocreatine, 4 Mg-ATP, 0.4 Na_2_-GTP and 0.3 % Biocytin (293-303 mOsm, pH 7.3). The membrane potential was amplified (Multiclamp 700B, Molecular Devices) and sampled at 20 kHz using WaveSurfer (wavesurfer.janelia.org). Membrane potentials were not corrected for the liquid junction potential. After the recording, the craniotomy was covered with Kwik-Cast (World Precision Instruments). Each animal was used for two or three recording sessions. Recordings were made 235 to 818 μm (521.6 ± 120.8 μm, mean ± s.d) below the pia. Brief current injections (-100 pA, 100 ms) were applied at the end of each trial to measure input resistance, series resistance, and membrane time constant ^73^.

For current injection experiments (Fig. 3), we partially compensated for series resistance and injected a ramping current until action potentials disappeared ^74,75^. Actual membrane potential was calculated post hoc based on injected current and series resistance. Mean membrane potential was −48.9 ± 3.4 mV (mean ± s.d., *n* = 10) without current injection. We injected −200 ± 153 pA, resulting in Vm = −60.9 ± 4.5 mV (mean ± s.d., *n* = 10). Series resistance did not change before and after current injections (*p* = 0.447, *ranksum* test, *n* = 10). In 5 / 10 recordings we were able to release current injections at the end of experiments to confirm that 1) membrane potential returned to spontaneous levels, and 2) neurons still produced action potentials.

### Whole-cell recording data analysis

Whole-cell recordings with more than 10 correct trials per direction (contra and ipsi trials) were analyzed (21.5 ± 8.2 correct trials per direction, mean ± s.d., *n* = 79). Performance during recording was 85.0 ± 11.8 % (mean ± s.d., *n* = 79). Only correct trials were analyzed. Cells recorded during the tactile task (n = 42) are from ref ^47^. (Supplementary Information Table 1). To measure membrane potential, spikes were clipped off (Fig. 2-4 and Extended Data Fig. 1-4). Neurons that differentiated correct trial-types during the delay epoch based on spike counts (100 ms averaging window) were deemed as “selective” (20 / 79 in ALM). To compute selectivity, we computed the difference in spike count or membrane potential between trial types (correct only) for each selective neuron (Fig. 2d, e).

To obtain spike-triggered median (STM) of Vm (Extended Data Fig. 1 and 2), we selected spikes that were not preceded by any other spikes in a 50 ms window. To obtain Vm-to-SR relationship, Vm and SR were averaged over a 50 ms sliding window. Mean SR was calculated for each step of Vm (1 mV step. Mean was defined for a step with more than 500 data points). For statistics of STM and Vm-to-SR relationship, we tested the null hypothesis that STM (or Vm-to-SR curves) in the pre-sample epoch and the delay epoch were identical. We performed hierarchical bootstrapping ^76–78^: we first randomly selected trials with replacement and then spikes within a trial with replacement. We measured the Euclidian distance of each bootstrapped STM (or Vm-to-SR curves) from the mean in the pre-sample epoch. The proportion of bootstrap trials with higher Euclidian distance in the delay epoch compared to that in the pre-sample epoch is shown as *p*-value. The results were robust to change in sliding bin size (50, 100 and 200 ms).

Delay activity of Vm or spike count (Fig. 3b and Extended Data Fig. 1b, 2b and 4c) was defined as the difference in Vm or spike count between the delay epoch and the pre-sample epoch. Spike bursts (Extended Data Fig. 1c, d and 2c, d) were detected as depolarization events 5 mV higher than spike threshold lasting longer than 20 ms. Vm autocorrelation and power spectral density were calculated after spike clipping (Extended Data Fig. 3). The time constant of membrane fluctuations was based on the autocorrelation function (Extended Data Fig. 3b) (time point when the function drops below 1/e).

To obtain across-trial fluctuations (Fig. 4 and Extended Data Fig. 5), we averaged Vm over 100 or 200 ms sliding window. We used first and third quartile difference (difference between 75 % point – 25 % point) or trimmed std (standard deviation after trimming off max and min data point) to calculate the across-trial fluctuations at each time point. This procedure removed the effects of a few outlier trials with spike bursts. For statistics, we performed hierarchical bootstrapping: first we randomly selected cells with replacement, and second randomly select trials within a cell with replacement.

### Extracellular electrophysiology

A small craniotomy (diameter, 0.5 mm) was made over the left ALM hemisphere one day prior to the recording session. Extracellular spikes were recorded using Janelia silicon probes with two shanks (250 μm between shanks) (Part# A2x32-8mm-25-250-165). The 64 channel voltage signals were multiplexed, recorded on a PCI6133 board (National instrument) and digitized at 400 KHz (14 bit). The signals were demultiplexed into 64 voltage traces sampled at 25 kHz and stored for offline analysis. One to five recording sessions were obtained per craniotomy. Recording depth (between 800 μm to 1100 μm) was inferred from manipulator readings (Supplementary Information Table 1). The craniotomy was filled with cortex buffer and the brain was not covered. The tissue was allowed to settle for at least five minutes before the recording started.

### Extracellular recording data analysis

The extracellular recording traces were band-pass filtered (300-6000 Hz). Events that exceeded an amplitude threshold (4 standard deviations above the background) were sorted using JRclust ^79^.

For the fixed delay auditory task with 2 s delay (Fig. 5, 6 and Extended Data Fig. 6, 9), in total 755 single units were recorded across 20 behavioral sessions from 6 animals (Supplementary Information Table 2). The same dataset was analyzed in ^47^. For the random delay auditory task (Fig. 7 and Extended Data Fig. 10), in total 302 single units were recorded across 11 behavioral sessions from 4 animals (Supplementary Information Table 3). For the calibration of photoinhibition without a behavioral task, 316 single units were recorded from 2 animals (Extended Data Fig. 8).

Spike widths were computed as the trough-to-peak interval in the mean spike waveform. The distribution of spike widths was bimodal (data not shown); units with width < 0.35 ms were defined as putative fast-spiking (FS) neurons (142 / 1373), and units with width > 0.5 ms were defined as putative pyramidal neurons (1202 / 1373). This classification was verified by optogenetic tagging of GABAergic neurons (data not shown) ^11^. Units with intermediate spike widths (0.35 - 0.5 ms, 29 / 1373) were excluded from our analyses.

Neurons with significant selectivity (*ranksum test* comparing spike counts in two correct trial types, *p* < 0.05) during the delay epochs were classified as selective cells. Selective cells were classified into “contra-preferring” versus “ipsi-preferring”, based on their total spike counts during the delay epoch. To compute selectivity, we took the firing rate difference between two correct trial types for each selective neuron. For the random delay task, selective neurons were defined based on the delay epoch of trials with 1.2 s delay (*ranksum test*, *p* < 0.05).

For the peri-stimulus time histograms (PSTHs; Fig. 6, 7), only correct trials were included. For the PSTHs and selectivity of random delay task (Fig. 7), only spikes before the go cue were pooled. Spikes were averaged over 100 ms with a 1 ms sliding window. Bootstrapping was used to estimate standard errors of the mean.

For Extended Data Fig. 9a, we compared spike rates of all unperturbed trials versus all perturbed trials using *t*-test. Cells with *p*-value lower than 0.05 was counted as significant cell at each time bin. For this plot, we did not correct for multiple comparisons.

For Extended Data Fig. 9c, d, we calculated selectivity for correct and incorrect trials based on the last 1 s of the delay epoch. We defined the polar coordinates *r* and *θ* as below:

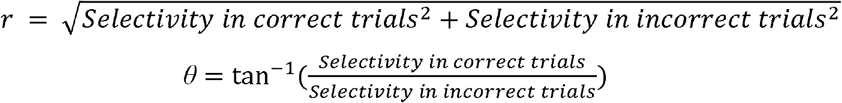

For Extended Data Fig. 9d, we pooled cells with *r* > 2 and *θ* in the range indicated in the figure. Many selective neurons in ALM have *θ* not at −45, indicating mixed coding ^47^.

To quantify the recovery time course of selectivity after photoinhibition (600 ms after the onset of photoinhibition, see main text), we looked for the first time bin when selectivity in photoinhibition trials reached 80 % of the control selectivity.

### Coding direction analysis

To calculate coding direction (CD) for a population of n simultaneously recorded neurons, we looked for an n × 1 vector maximally distinguishing two trial types (contra and ipsi trials), in the n dimensional activity space. For each neuron, we calculated average spike rates in the contra and ipsi trials separately (with 100 ms average window and 10 ms sliding step). *r*_contra_ and *r*_ipsi_ are n × 1 vectors of average spike rate at one time point. The difference in the mean response vector, *w_t_* = *r*_contra_ - *r*_ipsi_, showed high correlation during the delay epoch ^16,19^. We averaged *w_t_* during the last 600 ms of the delay epoch and normalized by its own norm to obtain the CD. We calculated CD based on randomly selected 20 % of unperturbed correct trials. To obtain trajectories along CD, we projected the spike rate in the remaining trials to the CD as an inner product.

Approximately half of selective cells show preparatory activity ^16^, which anticipates upcoming movements, regardless of behavioral outcome (correct or incorrect). To analyze CD trajectories in incorrect trials, for Fig. 6 and Extended Data Fig. 9, we only analyzed sessions with more than 5 preparatory neurons (11 / 20 sessions, see Supplementary Information Table 2 for trial number). Preparatory cells were defined as cells with *r* > 2 and 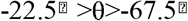 (Extended Data Fig. 8d, above). In Fig. 5, 7 and Extended Data Fig. 6, 10, all sessions were included as we only analyze correct trials (see Supplementary Information Table 2 for trial number). For Fig. 5, results were similar even when we only analyzed the sessions with more than 5 preparatory neurons (data not shown).

In Fig. 5, 6, projection to CD was normalized based on the value at the endpoint in each session. The distribution of endpoint values was bimodal (Fig. 5, 6). We calculated the median of each distribution (contra and ipsi) in unperturbed correct trials. The projection to CD was normalized by the difference of these two medians (ipsi peak at 0; contra peak at 1) in unperturbed correct trials. For Fig. 6e-h, and Extended Data Fig. 9e-g, we pooled trials from all sessions (with preparatory cells) after normalizing projections to CD in each session. Hierarchical bootstrapping was used to estimate standard errors of mean: first we randomly selected session number with replacement and, second randomly select trial number within a session with replacement.

Single trial projections to CD were noisy, because the point process noise scales with spike rate ^80^. To calculate across-trial fluctuations of projection to CD (Fig. 5d, e and Extended Data Fig. 6c) and drift (Fig. 5f and Extended Data Fig. 6f) for the fixed delay task, we rank-ordered and pooled trajectories per trial type based on the value 1.3 s before the go cue. For the random delay task (Fig. 7d and Extended Data Fig. 10j), we rank ordered and pooled trials based on the value at the onset of delay epoch, and analyze activities in the first 0.6 s of the delay epoch (we did not include trials with 0.3 s delay). Every 15 trials were pooled based on the rank order. Assuming that the point process noise is independent across trials, the averaging procedure reduced point process noise by 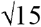. We calculated the standard deviation of the pooled trajectories at each time point. The first 100 ms was discarded, as the averaging window was 100 ms. The standard deviation was normalized by the value at 1.2 s before the go cue. As a control, we performed trial shuffling. We shuffled spike count among trials at each time bin independently. This procedure preserved the mean spike rate of each cell and CD but eliminates the time correlation within a trial. We performed the exact same procedure to calculate across-trial fluctuations of the shuffled data.

To calculate the drift of trajectories (Fig. 5f, 7d and Extended Data Fig. 6f), we analyzed the last 600 ms of the delay epoch. The drift of a trajectory at time *t* is defined as d*x*_*CD*_t_/dt = (*x*_*CD*_t+1_-*x*_*CD*_t-1_)/2, where *x_CD_t_* denotes projection to CD at time *t*. The time step was 10 ms. After pooling all time points and trajectories, the mean d*x*_*CD*_t_/dt was calculated for each step of *x_CD* (0.05 (a.u.) step. The mean was defined for a step with more than 50 data points. We only analyzed correct trials. In Fig. 5f and 7d both contra and ipsi trials were pooled. In Extended Data Fig. 6f, contra and ipsi trials were analyzed separately.

To decode future licking direction before the perturbation (Extended Data Fig. 9h-l), we analyzed the values of CD projection at the last bin 50 ms before the delay onset (*x*_*CD*_*−2.05*_). We first randomly selected 70 % of unperturbed correct trials. Next, we calculated the chance that animals lick right or left as a function of *x*_*CD*_*−2.05*_ (0.05 (a.u.) step) (Extended Data Fig. 9h). Based on this relationship we defined a decision boundary for lick right direction as the smallest *x*_*CD*_-2.05_ with probability of licking right that was higher than 70 %. The decision boundary for lick left direction was based on a similar procedure. We cross-validated these boundaries using the remaining 30 % of unperturbed correct trials and all of the unperturbed incorrect trials. Crossvalidated performance was 76.6 ± 16.7 % and 77.8 ± 15.8 % (mean ± s.e.m.) for lick right and left trials, respectively. We defined decision boundary and putative correct (or incorrect) trials independently for each session.

In extended Data Fig. 10e-i, to find modes explaining the remaining activity variance (RM, S1 and S2), we first found eigenvectors of the population activity matrix using singular value decomposition (SVD) at each time point and averaged over the last 600 ms of the delay epoch. The data for SVD at each time point was a n × trial-number matrix. We analyzed the first three eigenvectors from the SVD. All of these vectors were rotated using the Gram-Schmidt process to be orthogonal to CD and to each other. Since the projection to the first vector resulted in non-selective ramping activity in a fixed delay task (Extended Data Fig. 10e), we referred to this vector as a ramping mode (RM) ^16^. Projection to CD and RM were not normalized to the peak location in Extended Data Fig 10. To calculate the “selectivity explained”, we first calculated the total selectivity as a square sum of the selectivity across neurons (square sum of n × 1 vector). Then we calculated the square of selectivity of the projection along each mode, and divided it by the total selectivity. To calculate the “variance explained”, we first calculated the total variance as a square sum of the mean delay activity (difference between spike rate during the delay epoch and pre-sample epoch) across neurons in each trial type (contra and ipsi trial) (square sum of n × 2 matrix). Then we calculated the square sum of the mean delay activity in projection along each mode in each trial type. We divided this value by the total variance. For fixed delay tasks, we calculated selectivity and mean delay activity based on the last 600 ms of the delay epoch. For the random delay task, we calculated selectivity and mean delay activity based on the first 600 ms of the delay epoch.

### Network model

We built four firing rate models to simulate the average activity of neurons in ALM, (i) three discrete attractor network models (Fig. 6c right panel and Extended Data Fig. 7g-k, 11), and one integrator network model (Fig. 6c left panel and Extended Data Fig. 7b-f, l). We constructed attractor networks whose phase space contains two stable fixed points, corresponding to left and right licking, at the end of the delay epoch. Since we used the same laser power in both hemispheres to perturb neural activity in ALM, we first focused on modeling single-hemisphere dynamics, thereby ignoring inter-hemispheric interactions (Fig. 6c, Extended Data Fig. 7). To further account for the results of unilateral ALM perturbation experiments in Li et al, (2016) ^19^, we duplicated the one-hemisphere architecture shown in Fig. 6b, and linked the two identical modules together to portray both ALM hemispheres (Extended Data Fig. 11). Furthermore, we explored two potential mechanisms that may underlie slow ramping dynamics during the delay epoch: one where ramping was caused by a non-selective external input (Extended Data Fig. 11a-i), and one where ramping was a consequence of slow internal dynamics (Extended Data Fig. 11j-q). See Supplementary Information Table 4 for parameters used in the models described below.

### One-hemisphere discrete attractor model (Fig. 6c and Extended Data Fig. 7g-k)

We simulated the average activity of two excitatory populations in the left hemisphere, one selective to the right licking direction and one selective to the left licking direction, and one inhibitory population. The membrane current dynamics of excitatory population *i*, *h_i_*(*t*), was governed by the following nonlinear differential equation:

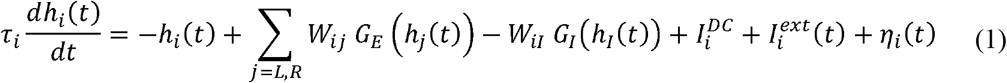

*W_ij_*denotes the synaptic strength between postsynaptic population *i* and presynaptic population *j*, labels *L* and *R* indicate the lick left and lick right selective excitatory populations,*I* denotes the inhibitory population, *τ_i_* is the integration time constant of the excitatory populations, 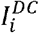 is the baseline input current, 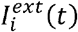 represents external input, *η_i_*(*t*) is a random Gaussian noise with zero mean and variance σ^2^. Since we did not explicitly model the dynamics of synaptic interactions, we included slow excitatory input such as NMDA in the excitatory time constant and set it to *τ_E_* = 100*ms* in all attractor models. The time-dependent external input 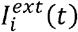 consisted
in population-specific selective input representing the sensory stimulus 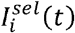(a 1 s long boxcar function exponentially filtered with time constant *τ_exp_* = 20*ms*) that was delivered during the sample epoch to either lick left or right excitatory neurons, depending on the trial type. The peak amplitude of 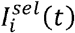 was drawn, across trials, from a Gaussian distribution of mean 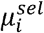 and standard deviation 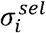. The transduction function *r_i_* = *G_E_*(*h_i_*), relating the firing *r_i_* rate to the current *h_i_*, is sigmoidal. We chose an easily interpretable parametrization of the transduction function, mimicking the effect of short-term synaptic plasticity ^81^:

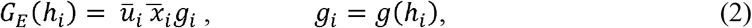

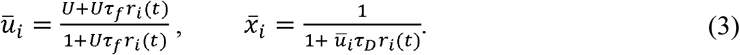

Variables *ū_j_* and 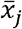 are steady-state synaptic nonlinearities resulting from, respectively, shortterm facilitation (STF) and depression (STD) of excitatory synapses; *U* is the synaptic release probability, *τ_f_* and *τ_D_* are the facilitation and depression recovery time constants (for their values see Supplementary Information Table). The activation function *g*(*x*) displays an exponential behavior around *x* = 0 and linear one for *x* ≳ *k*:

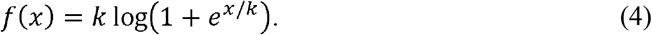

With these choices, the transduction function *r_t_* = *G_E_(h_i_*) exhibits a sigmoidal shape. This parameterization allowed us to make the basin of attraction of stable fixed points larger and more robust to noise. For the inhibitory population, we assumed instantaneous integration of the current received from the excitatory populations. We modeled the inhibitory transduction function *G_I_* as threshold-linear. The inhibitory firing rate can thus be written as:

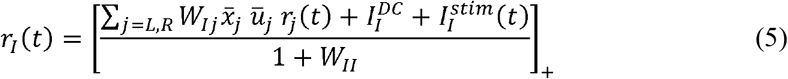

where 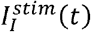 denotes the increase in the external input due to optogenetic photostimulation. 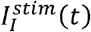 was modeled as a 600 ms long, exponentially filtered, positive input delivered at the end of the sample period. The value of the baseline input current 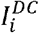 was such that *r_I_* was always greater than zero. For this reason, we replaced *G_I_*(*h_I_*(*t*)) in (1) with the argument of the threshold linear function in (5) to obtain:

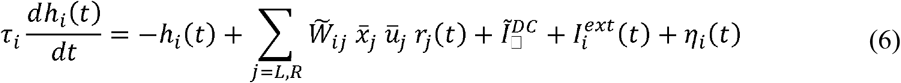

where now the synaptic strengths 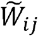 and the baseline input 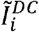 are rescaled according to:

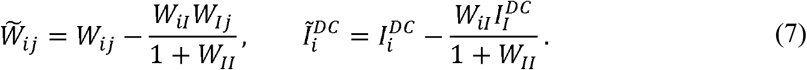

The synaptic weights were chosen such that the phase space contained two stable fixed points (Fig. 7f) during the delay epoch, corresponding to neural activity during contra and ipsi trials and one fixed point during the presample epoch, corresponding to baseline activity. The network switched between these two regimes 50 ms after the beginning of the sample epoch when the excitatory baseline currents 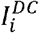 were significantly increased. The selective input current was drawn from the normal distribution 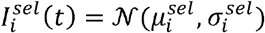 and was conveyed to lick right (left) population during sample epoch of contra (ipsi) trials.

### Two-hemisphere discrete attractor models (Extended Data Fig. 11)

To model ALM in both hemispheres, we linked together two identical one-hemisphere circuits (Extended Data Fig. 7a) via mutually excitatory coupling (*W_c_*) between lick left populations in one hemisphere and lick right in the other hemisphere. (Extended Data Fig. 11a). Each hemisphere, thus, contained one cell selective for licking right and one cell selective for licking left, both connected to an inhibitory cell. During the sample epoch the amplitude of the selective input, randomly varying from trial to trial 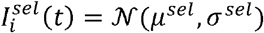 was delivered to lick left or right excitatory neurons of both hemispheres, depending on the trial type.

To reproduce the slow ramping of neural activity observed in the electrophysiological recordings (Fig. 6e), we considered two potential mechanisms in our simulations: i) a monotonically increasing nonselective external input *I^ramp^*(*t*); ii) the existence of a weak vector field along the direction of input integration, capable of slowing down the network dynamics during sample and delay epochs. In i), a nonselective input current *I^ramp^*(*t*), linearly ramping during the sample epoch and plateauing at the end of the delay epoch, was included in the external current 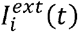 of all excitatory populations. We set the network parameters, including the peak amplitude of *I^ramp^*(*t*) such that the network would develop two stable fixed points during the delay period, corresponding to left and right licking directions. A single fixed point, corresponding to the baseline firing rate, was present all along the presample epoch. The nonselective ramping input destabilized the presample fixed point, while creating two stable fixed points during the sample epoch, corresponding to the left and right licking directions (Fig. 7f, and Extended Data Fig. 11a-i). Conversely, in ii) the network parameters were chosen so that the neural activity would slowly converge towards the fixed points (Extended Data Fig. 11j-q). In this configuration, which demanded substantial fine tuning, the phase space displayed two decision-related fixed points (left or right licking) during all epochs.

### Integrator model (Fig. 6c and Extended Data Fig. 7b-i, l)

Integrator dynamics was modeled using the negative derivative feedback mechanism ^23^. The firing rates of three populations (Extended Data Fig. 7a), were driven by recurrent synaptic inputs *s_ij_*(*t*):

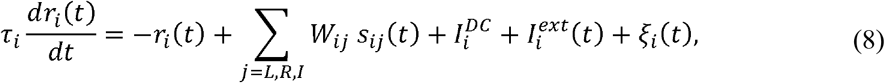

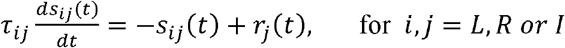

where *τ_ij_* are the recovery time constant of synaptic state variables connecting presynaptic population *j* to postsynaptic population *i*, the noise term *ξ_i_*(*t*) is a colored Gaussian noise with zero mean and two-point autocorrelation function 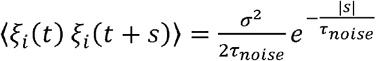 where *σ*^2^ the variance of the Gaussian distribution and *τ_noise_*is the decay constant of the autocorrelation function. The other variables in Equation (8) play the same role as in the discrete attractor models. To maintain persistent activity during the delay epoch, the conditions were *i*) balanced synaptic strengths and *ii*) time constants of positive feedback are longer than those associated with negative feedback (Supplementary Information Table 4). The linear system in (8) can thus display persistent activity lasting several seconds (Fig. 6c, left panel). During contra trials, we delivered a positive input, with the same temporal dynamics as in the discrete attractor model, to the lick right selective population, while a negative input was delivered during ipsi trials. To obtain the reduction in across-trial variability as in Fig. 5d, we ran an additional set of simulations where the variance of *ξ_i_*(*t*) decreased exponentially with time, starting from the end of the sample period (Extended Data Fig. 7I).

### Analysis of simulated activity

Numerical integration was performed using the Euler-Maruyama method for all attractor models and second-order Runge-Kutta method for the integrator model. To mimic the optogenetic activation of inhibitory neuron we chose, for each model, four values of 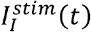 with increasing peak amplitude (see Supplementary Information Table). We simulated 1000 trials, of which 500 were contra and 500 ipsi trials, for each condition and model. All trajectories were smoothed using a 100 ms average window with 1 ms sliding step. The CD was computed using the same procedure adopted in the analysis of extracellular recordings (*r*_contra_ - *r*_ipsi_). Neuronal activity in each trial was then projected onto the CD to obtain activity traces and subsequently normalized using the distribution of endpoints in each session (see *Coding direction analysis* section). As with experimental data, we then analyzed correct and incorrect trials. Lick left trials were classified as correct if the neural activity of the lick left (lick right) selective population was higher than the neural activity of lick right selective neurons at the end of the delay period. In all attractor models, projected trajectories of trials classified as correct converged to the expected stable fixed point after bilateral or unilateral perturbation (Fig. 6c, Extended Data Fig. 7 and 11 all panels except 7d, i and 11d, i). Instead, trajectories of incorrect trials showed switching towards the wrong fixed point (Extended Data Fig. 7d and 11d). We calculated the drift of projected trajectories for each model using the last 1000 ms of the delay epoch. We applied the same method as in experimental data (Extended Data Fig. 7f, k and 11g, h, o, p). Fluctuations across trials were computed by taking the standard deviation across projected activity traces during correct trials, and normalizing them to their value at the beginning of the delay epoch (Extended Data Fig. 7e, j, l and 11f, n). Model performance was computed by dividing the number of correctly classified trials by the total number of trials in each condition (Extended Data Fig. 11i, q).

### Statistics and data

The sample sizes are similar to sample sizes used in the field. No statistical methods were used to determine sample size. We did not exclude any animal for data analysis. During experiments, trial types were randomly determined by a computer program. During spike sorting, experimenters cannot tell the trial type, so experimenters were blind to conditions. All comparisons using *signrank* and *ranksum-tests* were two-sided except Fig. 5e where we tested decrease in value from the baseline. All bootstrapping was done over 1,000 iterations. Data sets will be shared at CRCNS.ORG in the NWB format. For network models, Matlab code will be made available for download.

